# President Roosevelt’s lions reveal a century of population fragmentation in Africa’s largest carnivore

**DOI:** 10.1101/2025.02.26.640359

**Authors:** Ellie E. Armstrong, Caitlin Curry, Katherine A. Solari, Simon Morgan, Bruce D. Patterson, Bernard Agwanda, Lillian D. Parker, Nancy McInerney, Lauren E. Helgen, Kristofer M. Helgen, Craig Packer, Dmitri A. Petrov, Elizabeth A. Hadly, Jesús E. Maldonado, Robert C. Fleischer, Michael G. Campana

## Abstract

Over the past century, lion (*Panthera leo*) populations across Africa have experienced rapid and severe declines. Despite this, East Africa is considered a modern-day lion stronghold. Here, we use whole-genome sequencing of both recent and historical lion populations, primarily collected during the Smithsonian’s Roosevelt East African (1909–1911) and Rainey (1911–1912) Expeditions, to investigate changes in population structure, connectivity, and diversity over the last ∼100 years in East Africa. We find a clear signal of population fragmentation when comparing historical (1896-1946) and recent (1990–present) lion populations. Our analyses reveal genetic distinctions between remaining lion populations in Kenya and Tanzania and document loss of genetic diversity over time including reduced heterozygosity and the accumulation of runs of homozygosity. We detect a severe bottleneck in both Kenya and Tanzania approximately 25 generations ago, coinciding with a severe rinderpest outbreak in the region that is known to have decimated bovid and ultimately carnivore populations in the area. Nevertheless, modern lions in East Africa still exhibit overall high levels of diversity and low levels of inbreeding. Our results provide direct evidence of the effects of increasing habitat fragmentation and the significance of temporal data for contextualizing current patterns of population connectivity and diversity.

## Main

“But the lions now offer, and have always offered, the chief source of unpleasant excitement.” –Theodore Roosevelt, *African Game Trails*

The lion (*Panthera leo*) was historically one of the most successful and widespread large carnivores on the planet. Their range once extended from the Cape of Good Hope to the Mediterranean, from Senegal to Somalia, and from Greece and Yemen, to central India^1,2^. Despite their iconic status and previous dominance across these landscapes, today lions occur only in western India and Sub-Saharan Africa. African lion populations have declined dramatically, over the last 60 years, with an estimated 20,000-25,000 individuals remaining^3^. Recent rates of decline are especially alarming, with the number of lions decreasing by 30% over the past two decades (approximately three to four lion generations) and by 48.5% since 1980^4^.

At present, there are two designated subspecies of lion based primarily on their geographic distribution and genetic similarity: *P. leo leo*, which includes lions from Asia and northern, western, and central Africa, and *P. leo melanochaita*, which includes lions from eastern and southern Africa^5,6^. Although there has been some disagreement about the demographic history that has given rise to modern diversity patterns of lions in the west, central, and Asiatic lineages (see ^1,2,7,8^), East Africa is thought to be the origin of the modern lion lineage^9,10^. Lions from East Africa exhibit higher genetic diversity^11–14^ compared to the other lineages and are also genetically distinct from lions in the southern lineage, notwithstanding their current classification within the *P. leo melanochaita* subspecies^5^.

East Africa is considered to be a modern lion stronghold^4,15,16^ and contains a majority of the African continent’s largest lion populations (i.e. populations with >500 individuals)^15^. Despite this, some genetic studies of lions from within this region show signatures of increased population structure and fragmentation, as well as reduced gene flow, e.g. in Kenya^17^, whilst genomic data from Tanzania suggests that there is still high connectivity between populations^18^. Furthermore, microsatellite data reveal that population diversity and connectivity in modern lion populations (including those in East Africa) are significantly decreased compared to historic levels^14,19^. Interestingly, patterns of diversity loss observed in microsatellite data are not reflected in mitochondrial DNA (mtDNA) markers, possibly due to female site fidelity and male dispersal behavior in lions^14,19^. MtDNA and microsatellites are also expected to vary in their ability to uncover relationships between individuals and/or populations over different timescales, since microsatellites are expected to evolve faster than mtDNA. However, previous studies investigating diversity loss using historic samples in lions have relied only on mitochondrial and <20 microsatellite loci^14,19^, both of which are known to limit population genetic inference due to being maternally inherited (mitochondria), allelic drop out (microsatellites), and having overall less power compared to genomic-level data. Moreover, microsatellite genotypes have high inter-observer and inter-laboratory error and can be difficult to retrieve from museum specimens (e.g. ^20^).

Historical samples can provide incomparable insights into the past distribution and diversity of species^21^. However, due to storage conditions, collection strategies, contamination, and the gradual degradation of DNA over time, these samples are not easily used for high-throughput genetics studies^22–24^. Studies using ancient DNA (broadly defined as DNA derived from museum, archaeological, palaeontological, and other non-optimal specimens) to examine changes over time usually do so from a phylogenetic perspective (i.e. to identify the relationships between historical or extinct and modern taxa) with small sample sizes. This is generally due to both preservation issues and opportunistic historical sampling, and the difficulty of amassing and coordinating specimens from different collections. However, as lions have long been a desirable focus of big game hunters and naturalists, there are numerous historical specimens housed in museum collections^25^. These collections provide a window into the past diversity and distribution of lions especially over the duration of their 20th century decline.

In 1909, former U.S. President Theodore Roosevelt embarked on an expedition to East Africa to collect specimens for the new Natural History Museum at the Smithsonian Institution^26^. The journey is documented in Roosevelt’s book *African Game Trails*^27^. Over the course of the two-year excursion (1909-1910), approximately 11,400 animal and 10,000 plant specimens were collected from what was formerly British East Africa (modern Kenya and Uganda), the Belgian Congo (modern Democratic Republic of Congo, DRC), and Sudan (modern South Sudan). Shortly thereafter, Edmund Heller, a zoologist who accompanied the Roosevelt expedition, joined a second expedition (1911-1912) to East Africa (modern Kenya) led by American businessman Paul J. Rainey with the goal of collecting additional specimens for the Smithsonian National History Museum. Together, these collections present an unparalleled picture of East African fauna and flora at that time and also provide a rare opportunity to examine change over time at the population-level within species with numerous carefully preserved specimens from the same area. Furthermore, the sampling period roughly coincides with the onset of rapid defaunation in the region.

The signing of the General Act of Berlin (1885) ushered in a rapid influx of British, French, and German settlers within wildlife dominated landscapes in East Africa. The region was highly valued for its vast, fertile lands which were seen as ideal for crop production, and it was also considered an exotic and exciting travel destination for the world’s wealthy^29,30^. During this time, many African peoples were displaced from their lands while railroads and cities were rapidly constructed. Colonial disruption of both people and wildlife in the region expanded over the next century, with conflicts related to white settlement, both world wars, regional political and social upheaval, and independence movements. Growing human populations and the legacy of colonial inequalities in the region have led to indirect and direct conflicts between humans and wildlife that continue today. Impacts on lions include traditionally driven and retaliatory killings ^31–34^, modern sport hunting^35^, and major mortality events from disease, such as canine distemper virus and rinderpest^36–38^. Habitat loss, reduced habitat connectivity, and declining prey populations have also contributed to severe declines^39–42^.

Here, we aim to quantify changes in population structure and connectivity of lions in East Africa over the last century. We leverage the extensive collection of lions amassed during the Smithsonian-Roosevelt and Rainey expeditions from what is now modern-day Kenya to generate whole-genome resequencing data from 45 historic lions. We add these data to newly-generated whole-genome data for 94 recent lions from Kenya and Tanzania (Serengeti National Park), and combine them with all publicly available sequence data (both modern and historical) from East Africa (Kenya, Tanzania, Uganda, Somalia, Ethiopia) and surrounding countries (DRC, Malawi, Sudan). In total, we analyze 100 recent (1985-1999) and 102 historical (1896-1946) African lion genomes to provide an unprecedented population-level view of lion genetic diversity and connectivity over the last century.

## Results

### Library preparation and sequencing

For simplicity, we use current country names, rather than geographic designations used at the time of collection (e.g. we refer to British East Africa as Kenya for both historical and recent samples). We sequenced 94 recent (1998–present) Kenyan (n = 20) and Tanzanian (n = 74) lion genomes (Table S1 & S2; Figure 1) and reanalyzed six previously published recent lion genomes from East Africa^14,43,44^ and countries directly bordering East Africa. Altogether, we included 100 recent samples from Kenya (n = 21), Tanzania (n = 77), the DRC (n = 1), and Somalia (n = 1) (Figure 1). For details on sampling locations and national parks, see Table S1 and Supplementary Figure S1).

**Figure 1:**
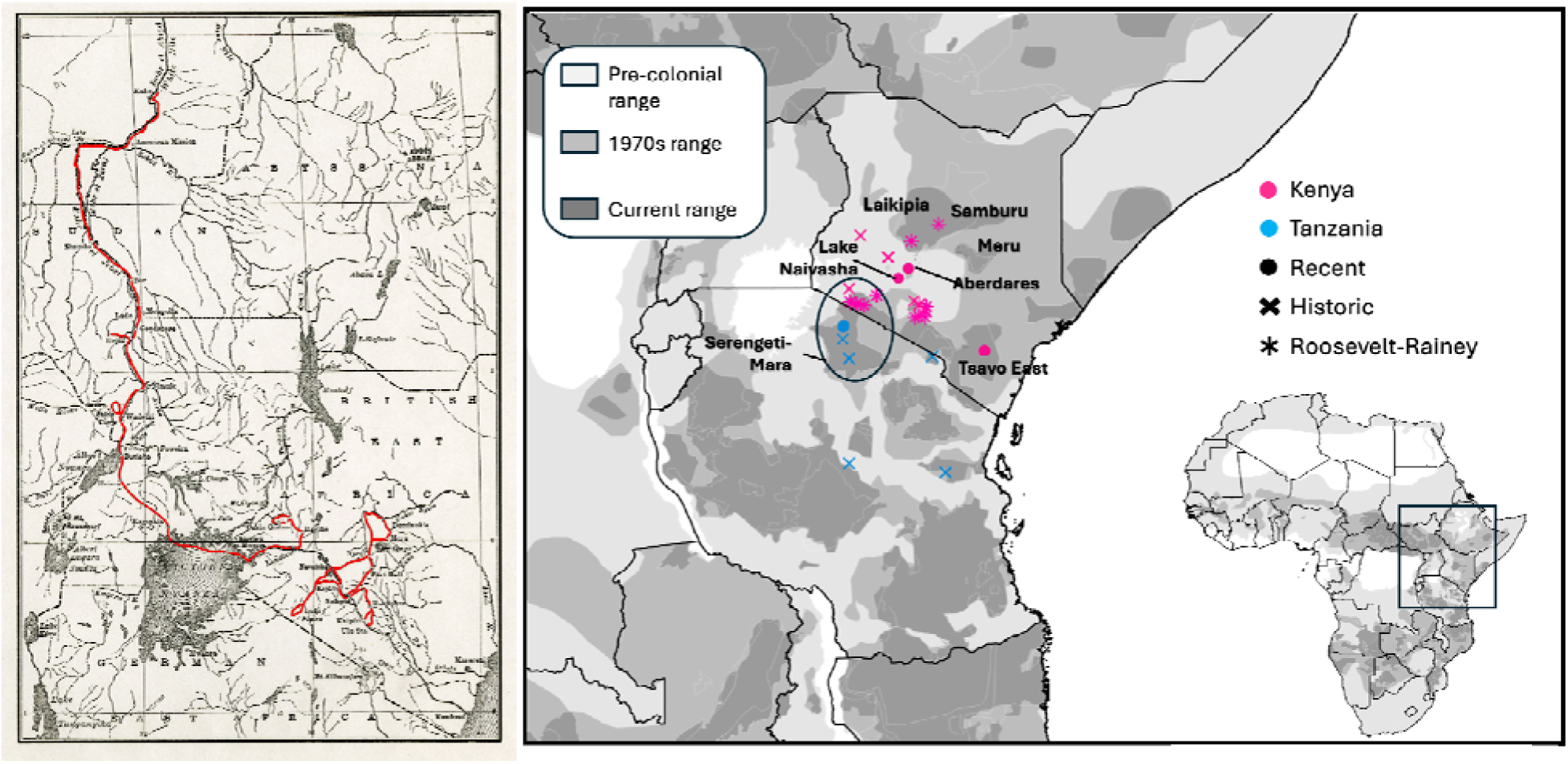
(Left) Map of Roosevelt-Africa Expedition from *African Game Trails* (Smithsonian Institution Archives). (Right) Map of approximate sampling location of individuals from Tanzania and Kenya included in this study, with notable regions identified. Range map data from^28^ and the IUCN RedList.

We sequenced 45 historical Kenya lions from the Smithsonian-Roosevelt and Rainey expeditions, (stored at the U.S. National Museum of Natural History, abbreviation USNM; Table S1; Table S2; note 45 of the 59 available samples were successfully prepared), and added 57 previously published^14,43^ historical genomes from East Africa and surrounding countries to our dataset. Our final dataset included 102 historical East African individuals (Kenya: n = 74; Tanzania: n = 24; Ethiopia: n = 1; Somalia: n = 1; Sudan: n = 1; Malawi: n = 1). Additionally, we deep-sequenced three historical Kenyan lions (USNM 161914, USNM 163108 and USNM 163109) to better investigate historical genetic diversity and inbreeding. We refer to the analyses including all samples sequenced at lower depth as ‘low-depth’ and those specific to the deep-sequenced libraries as ‘deep-sequenced’.

Recent genomes were sequenced to ∼4.8× after MapQ ≥ 30 filtering (mean ± SD: 4.77× ± 2.24×, range: 0.10×–11.95×; Table S3), while low-coverage historic genomes were sequenced to ∼0.2× (mean ± SD: 0.22× ± 1.15×, range: 0.00×–11.65×; Table S4). USNM 161914, USNM 163108 and USNM 163109 were further deep-sequenced to autosomal depths of 7.41×, 5.74× and 4.16× respectively on the Illumina Novaseq X 10B lane (Table S5).

### Dataset Generation

We generated three datasets encompassing different missingness thresholds to analyze the data (Table S4): a highest-coverage dataset including 53 historical individuals with sequences covering at least 10% of autosomal sites in the autosomal lion reference genome (≥ 6.2% after MapQ ≥ 30 filtering); a medium-coverage set including 62 historical individuals with sequences covering at least 5% of autosomal sites (3.00% after MapQ ≥ 30 filtering); and an ultra-low-coverage dataset including 74 historical individuals with sequences covering 1% of autosomal sites (0.60% after MapQ ≥ 30 filtering). All three datasets included all 100 recent individuals. For the three deep-sequenced individuals, we used only their initial shallow sequencing in the low-depth population analyses for data comparability.

For all three coverage cutoff datasets, we performed each analysis in two ways: 1) by including only transversion sites, as is typical in ancient DNA analyses to minimize the impact of cytosine deamination e.g.^45^; and 2) by including all variants (both transitions and transversions). At all coverage cutoffs where all variants were analyzed (1%, 5%, and 10% of sites covered), the transition/transversion ratio was ∼1.8, which is typical for mammals^46,47^. This indicates that our population-level data were not greatly impacted by excess artifactual transitions (Table S7). We ultimately excluded transitions from our final analyses as we observed inflated heterozygosities in the historical specimens when transitions were included (Supplementary Figure S13), though the results were overall similar across all analyses variations (Supplementary Data).

### Genetic sexing of historical samples

As a final quality-control step for our historical samples, we compared genetic sex assignments to sex assignments based on museum records. Of the 45 historical Roosevelt-Rainey samples, we assigned genetic sex to 33 individuals using the *R_x_* statistic (^48,49^; Table S1). Seven samples had fewer than 1,000 reads and five yielded ambiguous sex assignments. Of the 33 assignments, 30 (91%) matched the sex recorded in the museum catalog and three differed, all of which were recorded as male in museum records (USNM 182301, USNM 182305, USNM 182423). USNM 182423 was labeled as female on the skull, matching our genetic assignment, despite its catalog description listing it as male. Four genetic sex assignments were designated ‘tentative’ by the modified *R_x_* script (see methods); of these, three matched the museum records and one differed (USNM 182301). None of the three differing samples were included in the highest-coverage (10%) dataset. USNM 182301 was included in the medium (5%) and ultra-low coverage (1%) datasets, while USNM 182305 was only included in the ultra-low coverage dataset. USNM 182423 was discarded from all analyses due to its low sequencing depth (0.0012×; Table S3).

### East African Lion Population Structure

We assessed population structure on all three low-coverage datasets using principal component analysis (PCA) via PCAngsd^50^ and *K*-means clustering via ngsAdmix^51^ for *K* = 1–10. The results of all PCA and ngsAdmix analyses were concordant (Figure 2; Figure S2-S10). Principal component 1 separated recent Tanzanian lions (primarily from Serengeti National Park) from recent Kenyan lions (mostly from Tsavo East National Park). We found two recent Kenyan individuals (both from the Naivasha area) grouping with the majority of recent Tanzanian lions and one recent Tanzanian lion that was similar to extant lions from Tsavo. The inclusion of historical individuals revealed a north-south cline with historical individuals filling in the separation of recent populations in principal component space. Nevertheless, we found that historical Tanzanian and Kenyan individuals separated on the first principal component, but not to the degree of their recent counterparts. The second principal component separated three lions from the Aberdare Mountain Range (a highlands locality situated further to the northeast from our other Kenyan lion sample sites) from the remaining East African lions. Similarly, using a *K* = 2 model, ngsAdmix analysis found that historical samples were intermediate to their modern counterparts, with historical Tanzanian samples being more similar to recent Tanzanian individuals and historical Kenyan individuals being more similar to recent Kenyan individuals.

**Figure 2:**
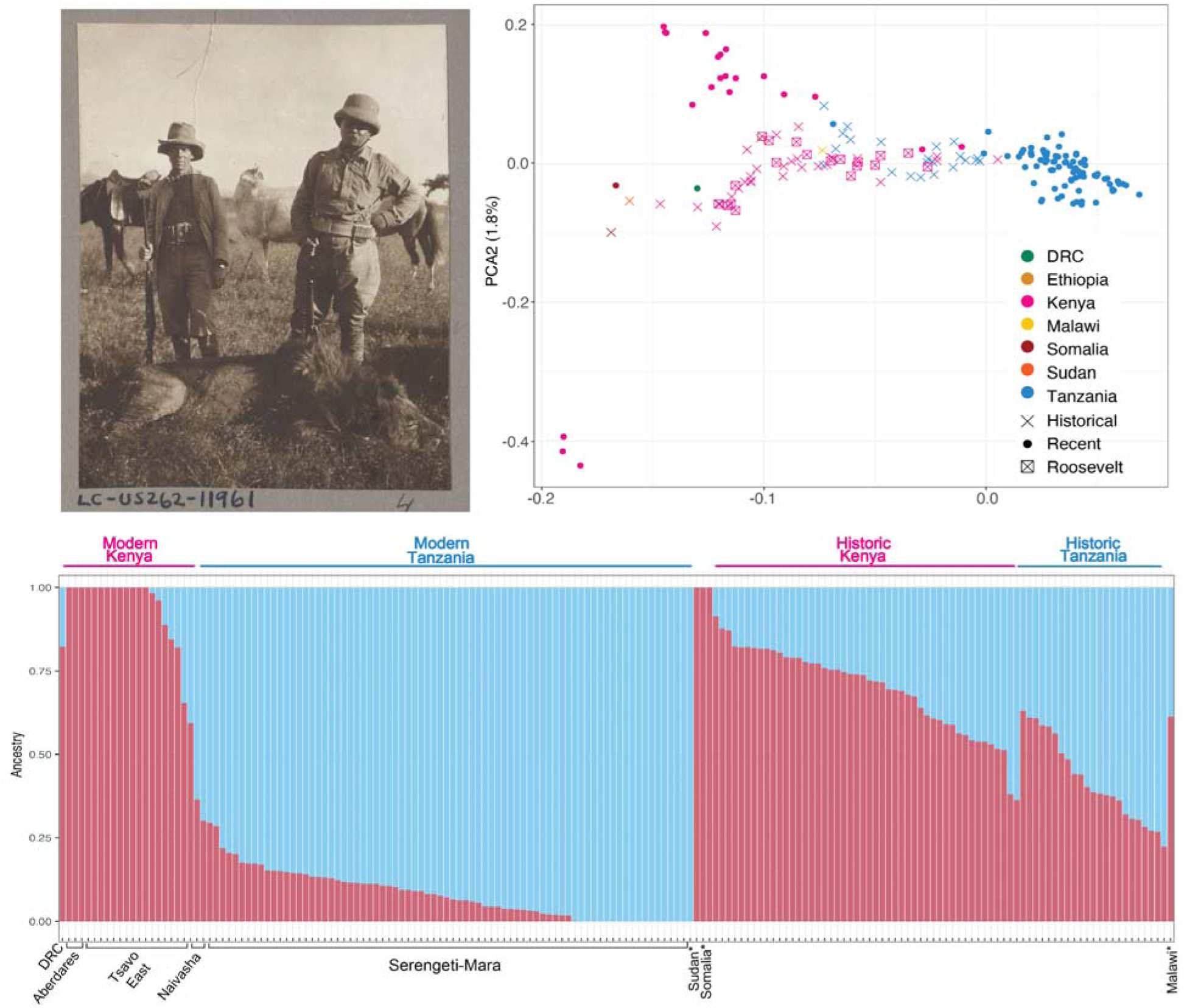
(Top left) Photograph by Kermit Roosevelt featuring President Roosevelt, Leslie Tarlton, and a male lion shot during the expedition (1909). (Top right) PCA of all samples with transitions removed. (Bottom) NGSAdmix results at *K* = 2 of all samples with transitions removed. Historical samples are noted either by a text descriptor (‘Historical’) or an asterisk (*).

Mitochondrial genome results clustered most Kenyan and Tanzanian individuals together (Figure 3), but reiterated the complexity of the mitochondrial patterns previously observed in lion populations across Africa^14^. Overall, we observed four major groups in the mitochondrial phylogeny, though the divergence between these groups is relatively shallow (note that branch labels are aligned for visibility and most average internal branches were between 0.001 and 0.003 or <150 substitutions between neighboring samples across the entire mitogenome). The first and second clades contain most of the samples from East Africa, as well as two samples from Zambia and one sample each from India and the DRC. These clades include most of the historical lions from East Africa. The historical samples tended to cluster together, but there were several of these clusters across the larger East African clade. The third clade included lions from Southern and Western Africa and the fourth clade included lions from all regions except Southern Africa. Overall, the mitochondrial phylogeny suggests a widespread sharing of lion mitochondrial haplotypes across Africa, likely due to the lack of strong barriers to lion movements, occasional long-distance female dispersal, and a large, panmictic historical population.

**Figure 3:**
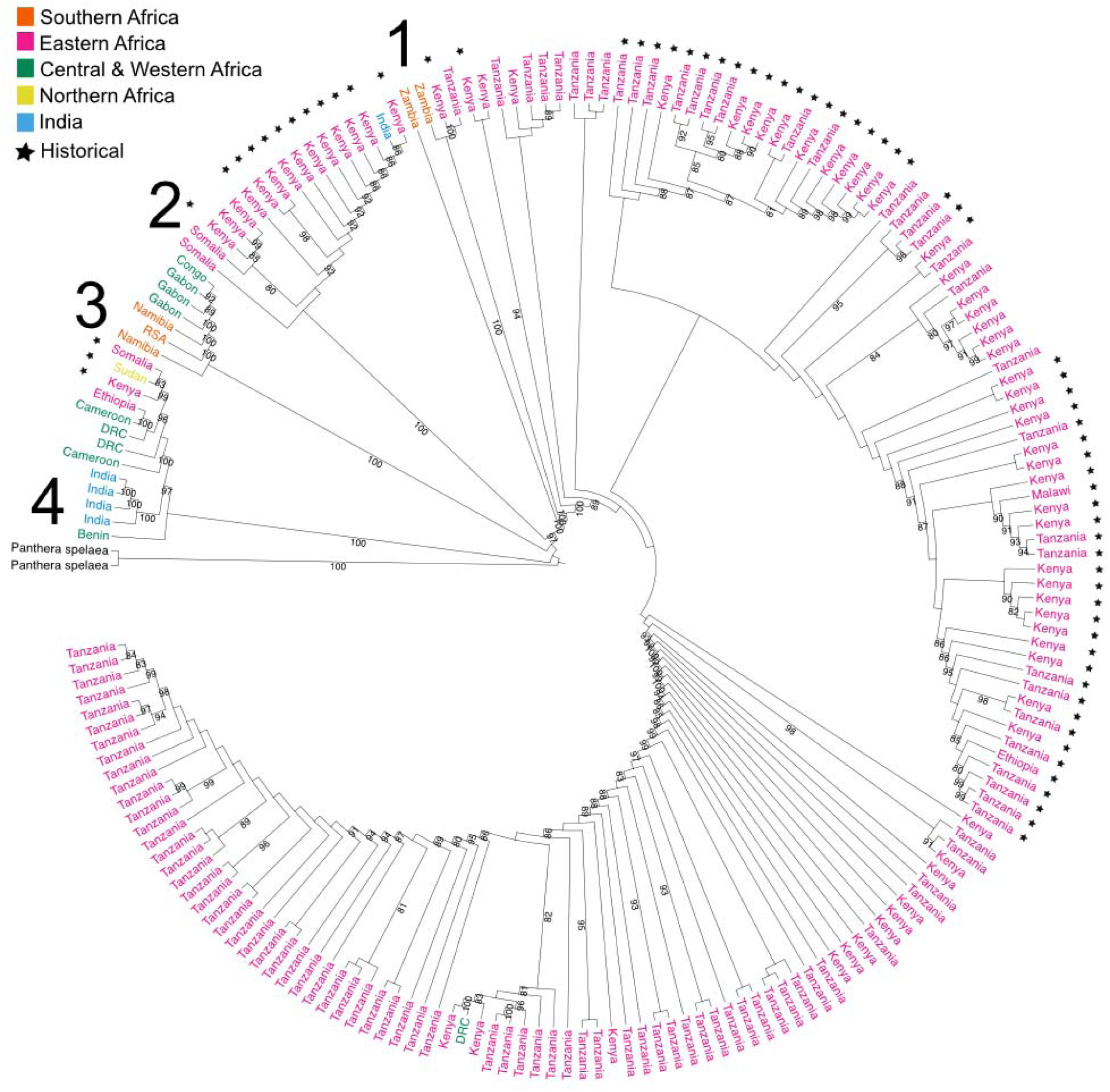
Mitochondrial phylogeny of historical and recent samples. Stars denote historical samples and bootstrap values >80 are displayed. Samples are colored by region (Eastern Africa, Southern Africa, Central and Western Africa, Northern Africa, and India) and the tree is rooted on samples of the extinct Eurasian cave lion (*Panthera spelaea*).

### East African Lion Diversity

In all low-coverage datasets, ngsF estimates of inbreeding showed that recent East African lions are significantly more inbred than their historical counterparts (10% coverage data transversions only, Wilcoxon test p << 0.0001, effect size r = 0.79; Supplementary Figure S11-12). The same pattern is observed when comparing recent Kenyan and Tanzanian lions (10% coverage data transversions only, Wilcoxon test p << 0.0001, effect size r = 0.81; Supplementary Figure S12) to historical lions from the same countries. Similarly, the three deep-sequenced historic lions from the Smithsonian-Roosevelt expedition are significantly more heterozygous than recent Kenyan lions (Wilcoxon test, *p* = 0.0091, effect size r = 0.52; Figure 4, Supplementary Figure S13-S14).

**Figure 4:**
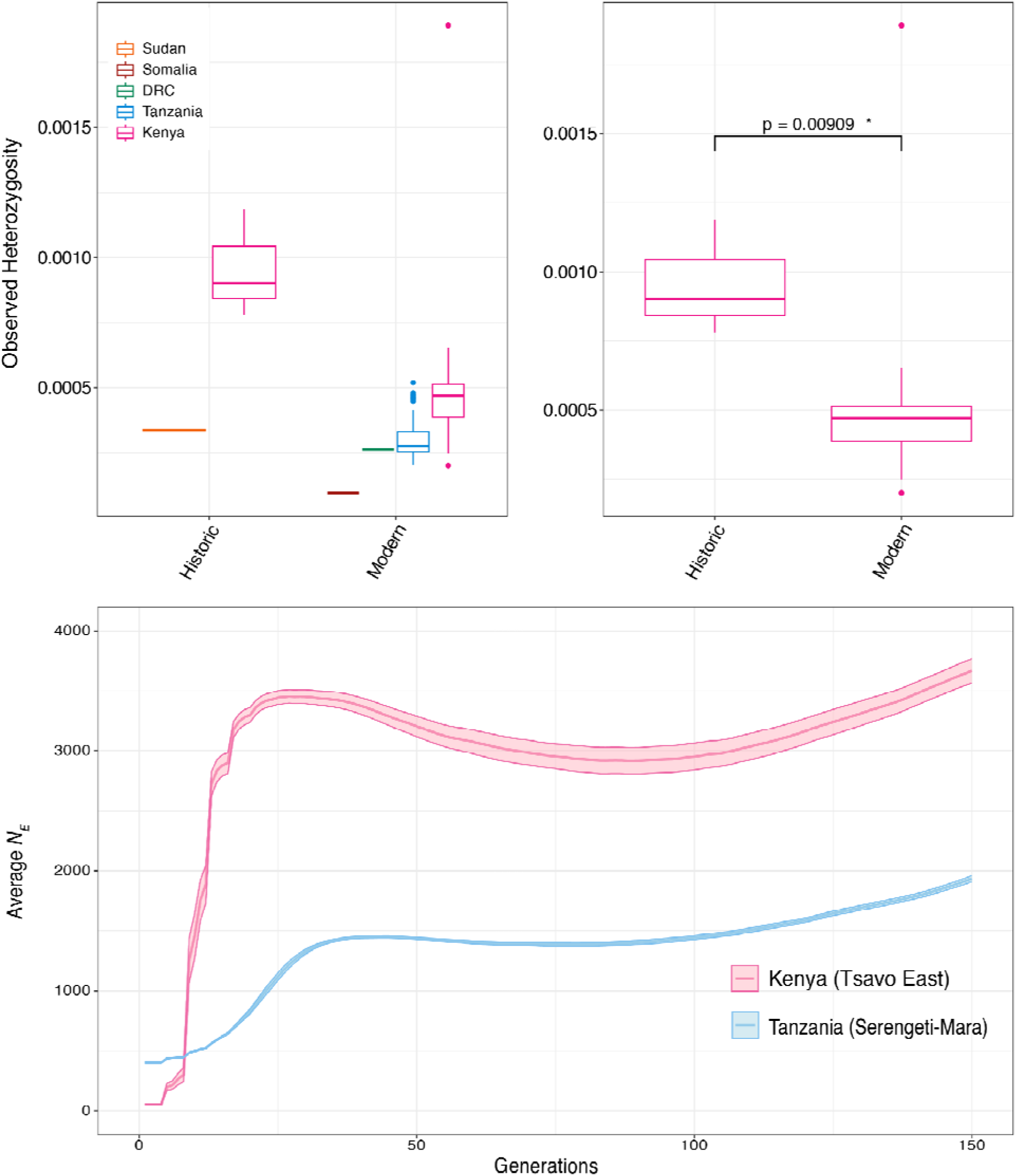
Heterozygosity of (top left) all historical and recent samples in East Africa and (top right) Kenya samples. Historic Ne estimates and 95% (shaded) confidence intervals for the last 150 generations for the Kenyan (Tsavo) and Tanzanian (Serengeti-Mara) populations.

We estimated runs of homozygosity (ROH) in samples with depths greater than 6× using ROHAN^52^ and found that overall lions have very few runs of homozygosity in their genome whether estimated with or without (Supplementary Figure S15-S17) transitions to accommodate the historical samples. Our results suggest that there has been an accumulation of ROH segments in recent lion populations (Supplementary Figure S17). As we only have three historical genomes over 6× depth, we take these results as suggestive rather than definitive. ROH is expected to increase when transitions are removed, which we observe in both recent and historical samples (Supplementary Table S15-S17). However, these runs are detected as short runs, rather than as long runs, when all sites are not considered. Furthermore, we show a comparable increase in the percentage of the genome in ROH when transitions are removed for both recent and historical samples (with transitions modern/historical: mean 0.38/0, without transitions recent/historic: mean 2.06/0.78), which suggests that our estimates are reliable.

### Demographic History

Using methods which rely on linkage disequilibrium, we estimated historical population size for the Serengeti-Mara (Tanzania) and Tsavo (Kenya) populations. Both populations showed a rapid population decline beginning approximately 25 generations ago (Figure 4). Historic effective population size in Kenya was much larger (∼3,500) than in Tanzania (∼1,500) prior to the bottleneck, which could also indicate that Tsavo was part of a larger or more diverse metapopulation prior to the bottleneck event or close to a suture zone where genetically differentiated groups were coming into contact. Current lion effective population sizes were estimated using current Ne at 202.39 (90% CI: 165.67-247.25) for the Serengeti-Mara and 11.60 (90% CI: 8.00-16.82) for Tsavo.

## Discussion

This study provides compelling evidence for significant loss of genomic diversity over the last 100 years in East African lion populations. Previous studies using historical samples have also shown signals of rapid diversity loss of lions associated with the period of European colonization in Sub-Saharan Africa over the past century. In Botswana, Dures et al.^19^ showed that recent samples had lower genetic diversity than historical samples and that mitochondrial haplotypes (using a 337 bp portion of the mitochondria) had been lost; due to data limitations, they did not investigate whether there were corresponding changes in population structure. Conversely, data from Curry et al.^14^ from across Africa showed no apparent loss of haplotypes using whole mitogenomes, but microsatellite data revealed apparent population panmixia for historic lions across their range. With the increased resolution of whole-genome sequencing, we show clear structure in East Africa in both historical and recent lion populations, but signals of substantial fragmentation in recent populations (Figure 2). Furthermore, we show genome-wide diversity loss and increased inbreeding in recent populations compared to historical populations (Figure 4, Figure S12 & S17). Our data underscore the utility of whole-genome sequencing in museomic studies comparing changes in genetic diversity through time as opposed to microsatellite or mitochondrial data given the inconsistencies in results between these marker sets from previous publications.

Our results provide insight into the status of recent East African lion populations. We showed that lions from the Aberdare Mountain Range (now Aberdare National Park) formed a distinct cluster from other Kenyan lion populations (Figure 2). Aberdare National Park contained one of the only montane populations of lions in East Africa and lies within a mountain range in the Great Rift Valley. Lions from this population were previously found to be genetically distinct from other lion populations^11^ and there has been some evidence to support a similar genetic pattern in other mammals (e.g. elephants^53^, African wild dogs^54^, sable antelope^55^, giraffes^56^). Historical samples in this study from the Laikipia region, even further north of the Aberdares, do not show the same distinctness as the Aberdares, suggesting the genetic uniqueness of Aberdares lions may be due to some level of upland isolation. Both the Laikipia and Aberdares region have seen significant declines in lions due to human-wildlife conflict^57,58^ and lions in Aberdares have been culled due to predation on other endangered ungulates^59^. Though we lack whole-genome recent samples from the Laikipia region, a previous study using a small custom SNP panel (n = 335 autosomal SNPs) suggested that Laikipia lions are not distinct from other nearby lion populations, possibly due to high levels of immigration^17^, though the resolution of this data may be limited due to the number of SNPs. The SNP panel also showed that the Solio Ranch population, which lies just to the east of Aberdares, was distinct from other lion populations, exhibited reduced genetic diversity^17^, and may be more closely related to the Aberdares population. The Solio Ranch population is fenced, possibly explaining its reduced diversity. We did not observe much lower values of genetic diversity (Table S1 and S8) in the individuals formerly occupying the Aberdares (n = 3), but additional samples would help determine the genetic distinctness of this population and whether lions in nearby areas may have a similar genetic makeup.

Interestingly, we observed the highest levels of recent diversity (H_O_; Figure 4) in Kenya, though the variance of these values in Kenya and Tanzania overlap. A recent census estimated the total lion population in Kenya at approximately 2,500 individuals ^60^). Comparatively, Tanzania supports approximately 10,000 lions^16^ and its populations together are purported to make up approximately half of all wild lion individuals. There has been significant concern with lion population declines in Kenya due to increasing habitat loss and fragmentation, whilst Tanzanian populations are considered large and relatively well-connected, however, a recent genetic analysis has suggested that lion populations in Tanzania are structured into northern, southern, and western groups ^61^. Though we observed one high-heterozygosity outlier in Kenya, the finding that individual heterozygosity is comparable (or higher) in Kenya compared to Tanzania is surprising. However, our demographic modeling shows higher historic population sizes in Kenya than in Tanzania, which explains the higher heterozygosity levels in Kenya. Previous work has shown that lions from northern Kenya (e.g. Meru, Samburu, and Maralel areas) and adjacent Ethiopia are genetically distinct from those in southern Kenya (e.g. Amboseli, Tsavo) and Tanzania^17^. Therefore, it is possible that Kenyan lion diversity has been maintained in part through translocations and/or migration of individuals across a genetically distinct suture zone present in equatorial Kenya. Admixture of these two distinct lineages could elevate genome-wide heterozygosity in Kenya beyond what modern population sizes alone suggest through the maintenance of divergent haplotypes encompassing multiple linked SNPs. Additionally, lion populations in East Africa appear to have undergone a severe decline in the 1890s due to a rinderpest (cattle plague) outbreak that is estimated to have reduced populations of bovids by 90% and killed countless additional wildlife indirectly through the loss of prey^62^. This signal is reflected in the demographic history as a bottleneck ∼25 generations ago in both Tanzania and Kenya, corroborating our results. More comprehensive sampling across Kenya and Tanzania will further contextualize some of these emerging patterns and the degree to which historic population size and/or admixture have shaped the genomic diversity in these stronghold lion regions.

Two recent Kenyan individuals (putatively from the Naivasha area) clustered with recent Tanzanian lions (which are primarily from the Serengeti; Figure 1). Naivasha is closer geographically to the Serengeti than the other modern Kenya samples sequenced here (Figure S1) and there may still be substantial connectivity between Naivasha and the Masai Mara/Serengeti ecosystem, as neither are technically fenced. An additional recent individual labeled as being from Tanzania (M_TAN_135; Table S1) from a previous study ^14^ also clustered nearer to recent Kenya individuals (Figure 1), but the exact collection point of this sample is unknown. More dense sampling of lions from their remaining habitat with precise locations will be necessary to clarify areas where gene flow is not occurring, compared to areas where there is connectivity or isolation by distance patterns. However, it is clear from our genetic comparisons that recent lion populations demonstrate more fragmentation, compared to historical populations from a century earlier. This confirms what might be expected given that lions no longer occur across much of their previous East African range, including in some areas sampled historically in our study.

We recognize several limitations of our study. First, although we analyzed many historical genomes, few are of high-coverage and/or depth. As sequencing costs decline, we anticipate the possibility of more comprehensive sequencing of museum specimens at the population-level will become more feasible. Encouragingly, many of the historical samples here had high endogenous content (Supplementary Table S3), so they can be revisited in the future with minimal impact on museum specimens. Furthermore, though locality data for the Roosevelt lions are generally well-documented, lack of exact georeference data for many historical and recent samples precluded finer-scale, spatially explicit analyses. This underscores the need for rigorous recording of georeference information for all ongoing lion sampling, and the importance of collaboration for contextualizing historical sampling information with local knowledge.

Over the last several centuries, anthropogenically-driven biodiversity loss has resulted in the extinction and local extirpation of hundreds of species ^63^. For large, wide-ranging mammals, remaining populations typically occupy less contiguous habitats than in the past, and illuminating corresponding patterns of genetic diversity change requires access to well-documented historical specimens stored in museum collections. Our findings, using this approach, demonstrate significant diversity loss over a relatively short span of time (∼100 years, or 20 lion generations given a generation time of five years) in an economically and ecologically valuable species, the lion in Africa. The onset of this rapid loss of diversity corresponds with intensifying colonization of the African continent at the turn of the 18th century amplified by subsequent impacts including increasing habitat fragmentation and human-wildlife conflict. Furthermore, the impact of 20th century management decisions, such as the building of fences and the relocation and translocation of lions between management areas (reviewed in ^17,64^), may influence interpretations of past genetic diversity patterns inferred from modern sampling alone. The use of historical samples provides the necessary context to illuminate genetic diversity patterns prior to 20th century alteration of habitats and disruption of lion dispersal behaviors This historical context may be useful to consider in defining lion management units and making future management and conservation decisions, including initiatives involving “rewilding” and translocations. For example, our findings confirm that early 20th century lion populations in East Africa were more connected than current populations, suggesting that ongoing translocation efforts in these populations are unlikely to be detrimental and may assist in maintaining and restoring historical patterns of gene flow.

Change-through-time comparisons of genetic diversity within single species, like the lion, provide one important way to examine the environmental impacts of wildlife decline and fragmentation. Future studies, similarly drawing on modern and historical samples in combination, might focus on other implications of recent change in African ecosystems, such as examining the genetic health of wildlife populations, or identifying historically shifting trophic interactions between species as environmental impacts have unfolded. Our current age of biodiversity decline, ongoing habitat modification, and shifting climates underscores the critical importance of utilizing museum collections as a tool for identifying and responding to the drivers and consequences of these rapid environmental changes.

## End Notes

## Acknowledgments

We thank K. Terio for assistance with accessing samples from the Serengeti Lion Project. We thank Susette Castañeda-Rico for laboratory assistance and Esther Langan for helping sample the National Museum of Natural History collections and Darrin Lunde for access to collections. Tuya Yokoyama helped prepare genomic libraries for the recent samples. E.E.A. was supported by the National Science Foundation Graduate Research Fellowship Program and a Smithsonian Short Term Visitor Award. Funding from this work was provided in part by the Smithsonian Institution, National Geographic Society (Grant No. 8846-10), Woodtiger Fund awarded to E.E.A. and M.G.C, a Realizing Environmental Innovation Program award through the Stanford Woods Institute for the Environment to D.A.P., as well as a Howard Hughes Medical Institute Professor award to E.A.H. We also thank Roland Kays and Hillary Young for feedback and assistance over the course of the project.

## Authors’ Contributions

E.E.A., K.M.H., J.E.M., R.C.F., and M.G.C. conceived the study and designed the experiments; B.D.P., C.P., and K.M.H. provided samples; M.G.C. sampled the museum specimens; E.E.A., L.D.P, N.M., and M.G.C. conducted the experiments; E.E.A., K.M.H., D.P., J.E.M., R.C.F, and M.G.C. secured the funding; E.E.A. and M.G.C. analyzed the data; E.E.A and M.G.C. wrote the paper; all authors revised and approved the manuscript.

## Methods

### Recent specimen extraction, library preparation, and sequencing

We obtained recent (1990s) lion specimens stored at the Field Museum of Natural History (FMNH; Chicago, USA; n = 20) and Lincoln Park Zoo (Chicago, USA; n = 74) frozen tissue collections (Table S1). The samples from the Field Museum were collected for previous studies and stored at the Field Museum for long-term storage. Field Museum samples were previously collected for a study used to examine population genetic structure of lions using mitochondrial and microsatellite loci^13^. These samples originate from individuals euthanized by wildlife authorities as problem animals within Aberdares and Tsavo East and Tsavo West National Parks in Kenya; vouchers for many of these were subsequently deposited in the National Museums of Kenya, Nairobi. Additional details on sample collection can be found in^13^. Recent Tanzanian samples were collected by C.P. as part of the Serengeti lion project. These samples have been previously used for genetic studies (see e.g. ^38,65^) and stored by Lincoln Park Zoo.

DNA extracts were quantified using a Qubit v2.0 (Thermo Fisher Scientific, Waltham, MA, USA) fluorometer using the Qubit High Sensitivity dsDNA Assay with 2 μl of DNA input. Approximately 2.5 ng from each were used as input material for library preparation. A low-input library preparation method using a modified Nextera protocol was used^66^ in order to preserve as much of the original material as possible, while still obtaining full genomic coverage. Libraries were sequenced on an Illumina HiSeq X Ten with 2 × 150 bp paired-end reads at Admera Health (South Plainfield, New Jersey).

### Recent data mapping for population structure and diversity analyses

Using a previously published Nextflow pipeline^67^; https://github.com/campanam/Elephants; pipeline version 0.2.1), paired sequences were aligned to the lion mappable reference genome (which includes a single haplotype in addition to the X and Y chromosome; ^68^ using BWA-MEM v0.7.17 under default parameters ^69^. Alignments were converted to BAM format (view command with options ‘-b -F 4’) and co-ordinate sorted using SAMtools 1.18 ^70^. Indels were left-aligned using the Genome Analysis Toolkit (GATK) 4.4.0.0 LeftAlignIndel command under default parameters ^71^. PCR and optical duplicates were marked using SAMtools. Multiple libraries from the same individual were then merged using SAMtools and indel-realignment and duplicate marking were repeated as above.

### Historic specimen extraction, library preparation, and sequencing

We sampled adherent soft tissue and previously damaged bone from 59 museum lion specimens (Table S1) collected during the 1909–1911 Smithsonian-Roosevelt East African 1911–1912 Rainey expeditions and housed at the National Museum of Natural History, Smithsonian Institution (Washington, DC, USA). Museum tissues were extracted using a previously published phenol-chloroform protocol^72,73^ in the ancient DNA laboratory at the Center for Conservation Genomics, Smithsonian’s National Zoological Park and Conservation Biology Institute under stringent anti-contamination procedures appropriate for ancient DNA research^74–76^. Extracted DNAs were built into double-indexed double-stranded DNA libraries using KAPA Library Preparation Kits – Illumina (Kapa Biosystems, Wilmington, MA, USA) and iNext adapters^77^ following^78^ or the BEST 2.0 single-tube library preparation protocol ^79^ with the modifications described in ^80^. Samples from five individuals were built into double-indexed single-stranded libraries using the Santa Cruz reaction^81^. Libraries were sequenced on Illumina (Illumina, Inc., San Diego, CA, USA) platforms at Admera Health (South Plainfield, NJ, USA) and the University of California Berkeley DNA Sequencing Facility (Berkeley, CA, USA) using 2 × 150 bp paired-end reads or 100 bp single-end reads (Table S2). Of the 59 individuals sampled, 45 individuals produced quantifiable libraries and were used for sequencing (Table S2). Additionally, three of the Santa Cruz reaction libraries (corresponding to individuals USNM 161914, USNM 163108 and USNM 163109) were further deep-sequenced to an additional ∼6× on a Novaseq X 10B lane at Admera Health.

### Historical data mapping

Historical data were processed using a custom Nextflow^82^ pipeline. Reads were first trimmed using AdapterRemoval2 v2.3.3^83^ retaining reads at least 30 bp after trimming (options ‘--gzip --minlength 30’). For paired-end reads, we collapsed overlapping reads (option ‘--collapse’). We discarded uncollapsed read pairs and treated collapsed reads as single-end reads for downstream alignment.

We aligned the trimmed and collapsed reads to the lion mappable genome^68^ using BWA-Aln 0.7.17^84^ with the seed disabled (option ‘-l 1024’ ^85^) and sorted the alignment by coordinate using SAMtools 1.18^70^) *sort*. We then indexed the alignments using Picard 3.1.0 (https://broadinstitute.github.io/picard) and left-aligned indels using GATK 4.4.0.0 LeftAlignIndels. We then marked PCR and optical duplicates using SAMtools *markdup*. For each alignment, we then profiled ancient DNA damage using DamageProfiler 1.1^86^. After damage profiling, we merged data for individuals sequenced across multiple libraries and lanes using SAMtools merge and then reperformed indel left-alignment and duplicate marking as above. Finally, to minimize the impact of cytosine deamination artifacts, we trimmed 2 bp from each of the aligned molecules using BamUtil 1.0.15 *trimBam*^87^. We then calculated alignment depth, breadth of coverage, and flag statistics for each sample using SAMtools (depth, coverage, and flagstat commands respectively).

### Generation of genotype likelihoods and site-filtering

For each low-coverage dataset, we identified variants and calculated genotype likelihoods using ANGSD 0.941 (Analysis of Next Generation Sequencing^88^). We employed a genotype-likelihood approach because the difference of sample depths between the recent (∼4.8×) and historic samples (∼0.2×) and overall low depth (<12× per individual) is typically problematic for hard-calling variants^88^. ANGSD parameter settings were derived from^89^ (options ‘-doMajorMinor 1 -GL 2 -doMaf 2 -doGeno 7 -SNP_pval 1e-6 -doPost 1 -doCounts 1 -dumpCounts 4 -dosnpstat 1 -doHWE 1 -HWE_pval 1e-2 -minMaf 0.01 -P 10 -doSaf 1 -underFlowProtect 1 -minMapQ 30 -minQ 20’). We output genotype likelihoods in both beagle genotype likelihood format (option ‘-doGlf 2’) and beagle binary format (option ‘-doGlf 3’) for downstream analyses. We polarized the site frequency spectra using the lion reference genome (option ‘-anc <reference sequence>’) and restricted our analyses to autosomes (option ‘-rf <autosome list>’). We also reperformed these analyses including transversions only (option ‘-rmTrans 1’ and omitting incompatible option ‘-doSaf 1’).

### Genetic sexing of historical samples

Using SAMtools 1.18, we removed sequence alignments with mapping quality (MapQ) < 30 and recalculated genomic coverage after their removal. We discarded individuals with fewer than 1000 aligned reads after filtering. We then assigned genetic sex using the *R_x_*statistic^48^ as modified for lions following^49^.

### Diversity, kinship and inbreeding

For each low-depth ANGSD dataset, we calculated kinship and inbreeding coefficients using ngsRelateV2 under default settings^90^. We identified close kin (approximately third-degree) using *r*_ab_ > 0.10. We note that kinship and relatedness coefficients are frequently significantly overestimated when analyzing data with sequencing depth < 0.1×^91^, so we treat these results with caution. We also calculated inbreeding coefficients using ngsF 1.2-STD under default settings^92^.

We evaluated individual diversity by estimating heterozygosity per individual. We first generated a folded site frequency spectrum (SFS) using ANGSD v0.9.3.1 using the same flags for both the recent and historic samples which included: “-fold 1 -dosaf 1 -GL 2 -C 50 -minq 20 -minmapq 30”. We only considered autosomal regions and the reference genome was provided for both the ‘-ref’ and the ‘-anc’ flag in order to fold the SFS. Using the same flags as listed before, we also generated a second folded SFS with the ‘-noTrans’ flag, which removes transitions from consideration. Subsequently, *realSFS* from the program ngsTools was used to summarize the heterozygous and homozygous sites from the folded SFS. Observed heterozygosity was computed as the number of heterozygous sites over the total number of sites considered.

We used ROHan^52^ to estimate runs of homozygosity in recent and historic genomes with an average depth of at least 6×. We first estimated the transition/transversion ratio (Ti/Tv) in our samples by examining the number of variant sites that were removed by the ‘-noTrans’ flag in previous analyses (see Table 1). For the recent samples, we ran ROHan using the suggested parameters, including ‘-rohmu 2e-5’ and the calculated Ti/Tv of 1.8 with the flag ‘-tstv 1.8’. Autosomes IDs were provided using the ‘-auto’ flag. For historical samples, we tested the *bam2prof* module to explore damage profiles in our samples by increasing the minimum quality cutoff. We subsequently ran analyses with the following configurations: all sites for recent and historic samples with no damage profiles included; transversions only for recent and historic samples with no damage profiles included; all sites for recent and historic samples with damage profiles included; and transversions only for recent and historic samples with damage profiles included. Damage profiles provided utilized the ‘-l’ and ‘-minq’ flags set at 30, for a length cutoff and minimum quality cutoff score of 30, respectively.

### Population structure and connectivity

For each low-depth ANGSD dataset, we performed principal component analysis (PCA) using PCAngsd 1.21^50^ and assessed population structure using ngsAdmix^51^. For PCA, we modeled 4 eigenvectors and performed up to 1000 iterations to converge (options ‘--n_eig 4 --iter 1000’). For ngsAdmix, we assessed *K* values between 1 and 10 and performed up to 10,000 iterations (option ‘-maxiter 10000’). As a conservative confirmation of the observed spatial structure, we reperformed PCAngsd analyses after removing previously identified close-kin. The all-sites kin-pruned PCAngsd runs did not converge in 1,000 iterations, so these were rerun with up to 10,000 iterations for convergence (option ‘--iter 10000’). The all-sites kin-pruned 5% and 1% datasets did not converge within 10,000 iterations, so we interpret these results with caution. PCA and ngsAdmix results were plotted in R 4.2.1 using ggplot2 3.5.0^93^, patchwork 1.2.0^94^, and dplyr 1.0.10^95^.

### Mitochondrial DNA

Sequence reads were mapped to a verified lion mitochondrial genome sequence used in previous studies (KP001505.1^96^), in addition to a nuclear mitochondrial (numt) sequence from a tiger (DQ151551.1^97^). The tiger numt sequence was included in order to prevent reads from numt regions from mapping through competitive mapping. Besides changing the reference sequences, historical and modern samples used the same alignment pipeline as described above. We estimated depth and coverage of the mitochondrial genomes using samtools *coverage*. We screened all of the mitochondrial genomes for depth (the average number of reads covering each base) and coverage (the average number of bases covered). Only samples which had ≥50% of the mitochondrial genome covered and a mean depth of at least 10× were included for further analysis. Remaining mitochondrial genomes were aligned using mafft v7.520 ^98^ and subsequently trimmed using trimal v1.4.rev22 ^99^. We performed model selection using IQ-TREE v2.3.4 ^100^ by providing the trimmed alignment with flag ‘-m MFP’. We then ran IQ-TREE using the best model (-m K3Pu+F+I+R3) with 1000 bootstraps (-B 1000). The consensus tree was plotted using iTOL ^101^.

### Recent variant calling for demographic modelling

Sequence data was mapped using NVIDIA Parabricks v4.1.1.1 *fq2bam* with the provided reference and default parameters ^102^. We estimated depth and other mapping statistics using Mosdepth v0.3.3 ^103^ and SAMtools *flagstat,* respectively. Resulting BAM files were individually run through Parabricks *HaplotypeCaller* with the ‘--gvcf’ flag in order to emit both variant and invariant site calls. Subsequently, GATK v4.1.4.1 *GenomicsDBImport* was used to import the single sample VCFs prior to joint genotyping. Autosomal chromosomes were provided as intervals, excluding sex chromosomes and unlinked scaffolds for downstream analyses. Finally, we ran Parabricks *GenotypeGVCFs* VCF files. Files were combined using BCFtools v1.16 *concat* to create a single, autosomal VCF.

Samples with an average depth of less than 4× and more than 15% missing data were removed from the VCF using BCFtools *view*. VCF files were subsequently restricted to biallelic SNPs using BCFtools *view* using the flags ‘--min-alleles 2’, ‘--max-alleles 2’, ‘--exclude-types indels’. In addition, we excluded sites to the mappable genome (removal of low mappability and repetitive sites). As closely related individuals can skew demographic inference, we tested for relatedness using VCFtools ‘--relatedness’ and removed individuals that were 2nd degree relatives or closer. This resulted in six individuals from the Kenya Tsavo population and 44 individuals from the Serengeti-Mara population in Tanzania. We then restricted this file to sites with a minor allele count of one and a minor allele frequency of 0.1 using VCFtools. These were used as input for demographic modeling.

### Modern demographic history

We investigated recent demography using patterns of linkage disequilibrium as employed by GONE ^104^. We modified several default input parameters, including assuming a constant rate of recombination of 2 cM Mb^-1^ ^105^, with 500 replicates, and the parameter *hc* was set to 0.02. We ran this 500 independent times to obtain confidence intervals for *N_e_*. We subsequently estimated modern *N_e_* using currentNe ^106^ with default parameters on all 18 autosomal chromosomes.

## Data Availability

The sequencing reads generated during the current study are available in the Sequence Read Archive under Bioproject PRJNA1178729. The lion mappable genome described in^68^ and analyzed during the current study is available in the Smithsonian Figshare repository under doi: 10.25573/data.25374244, https://doi.org/10.25573/data.25374244.

## Code Availability

The historical DNA analysis Nextflow pipeline described in this manuscript is available in the GitHub repository, https://github.com/campanam/MuseumSpecimens. The modified sexing script (lion-rx.r) is available in the MuseumSpecimens GitHub repository. The previously published mapping and alignment pipeline used for the extant individuals is available in the GitHub repository, https://github.com/campanam/Elephants^67^.

