## Supplementary Figures for "President Roosevelt’s lions reveal a century of population fragmentation in Africa’s largest carnivore"

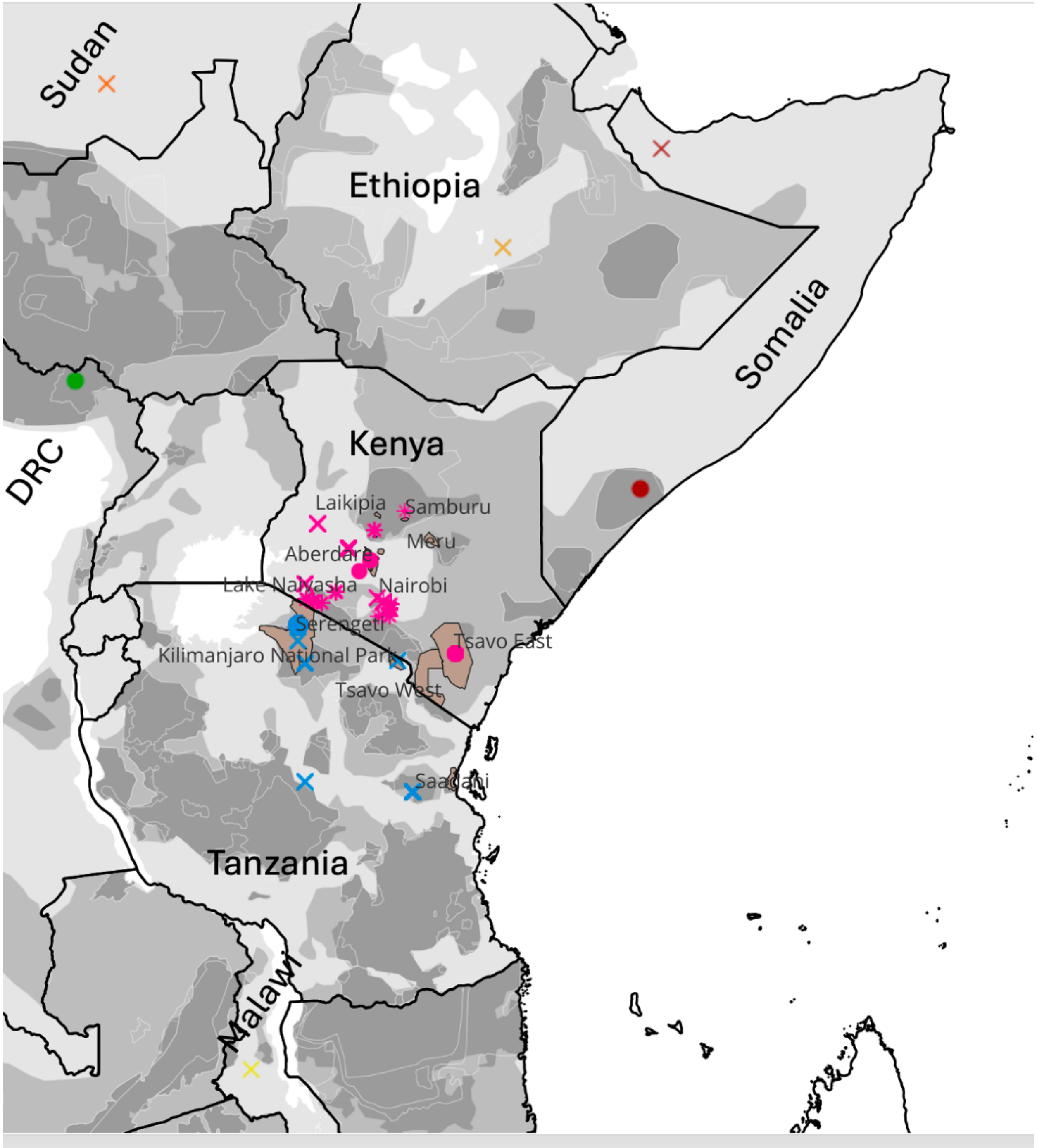


**Supplementary Figure S1:** Map of samples with relevant National Parks and Natural Reserves labeled.

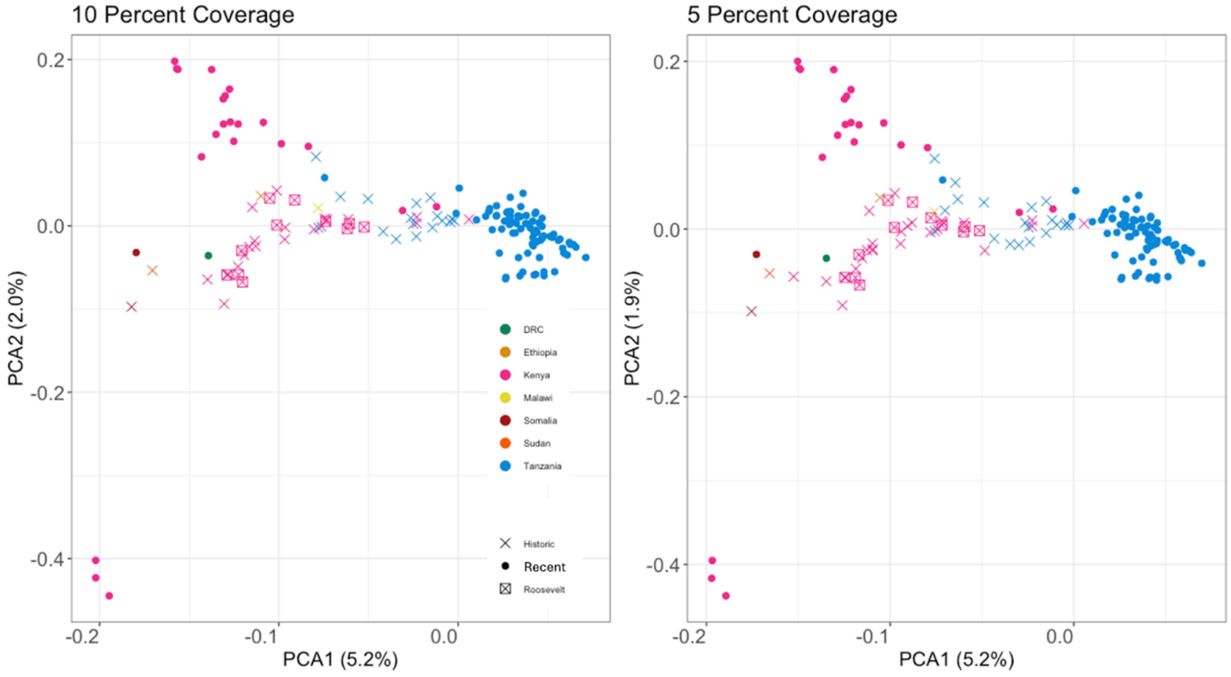


**Supplementary Figure S2**: PCA of recent and historic data with transitions removed across varying coverage cutoffs.

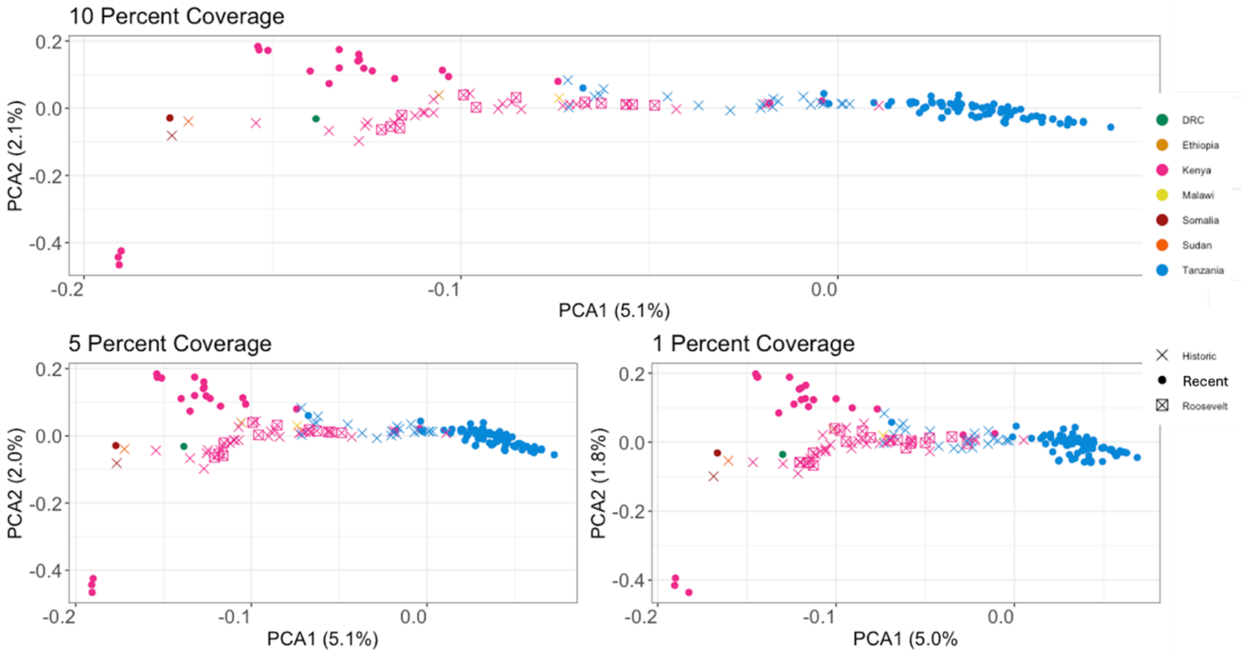

**Supplementary Figure S3**: PCA of all data (transitions included) across varying coverage cutoffs.

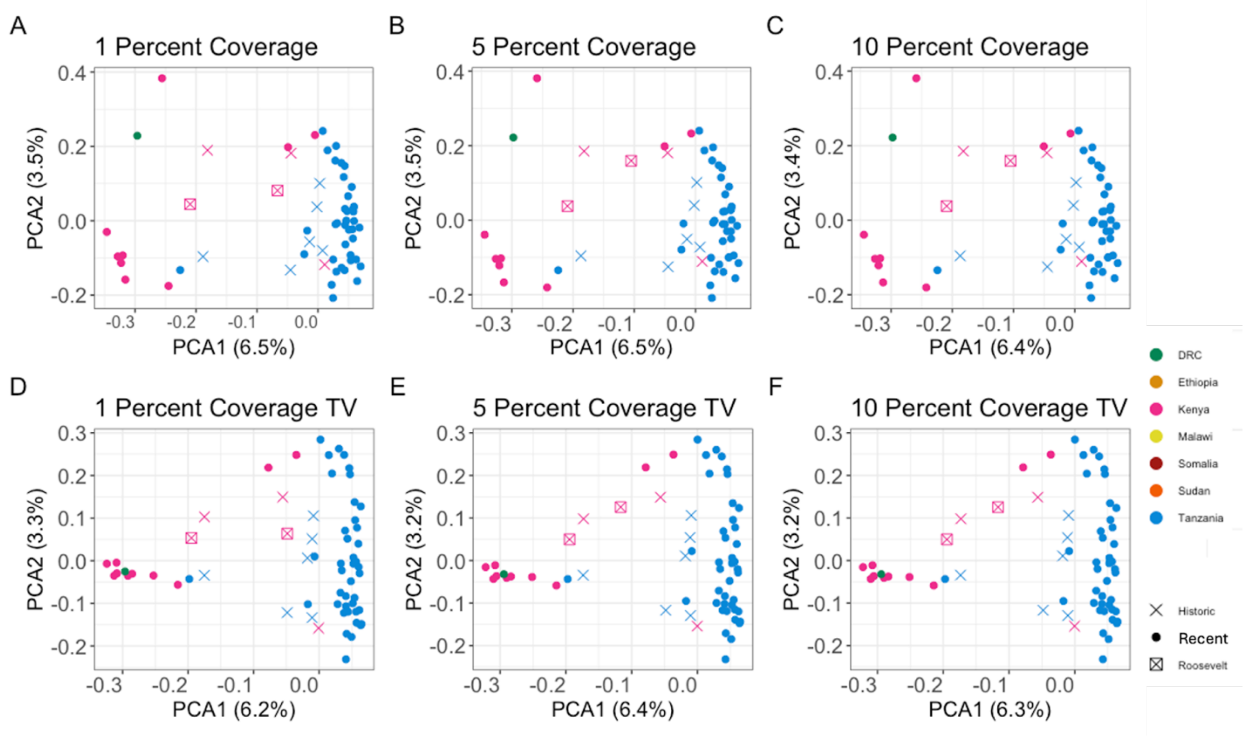


**Supplementary Figure S4**: PCA with relatives (3rd degree or higher) removed. With all sites (A-C) across varying coverage cutoffs. With transitions removed (D-F) across varying coverage cutoffs.

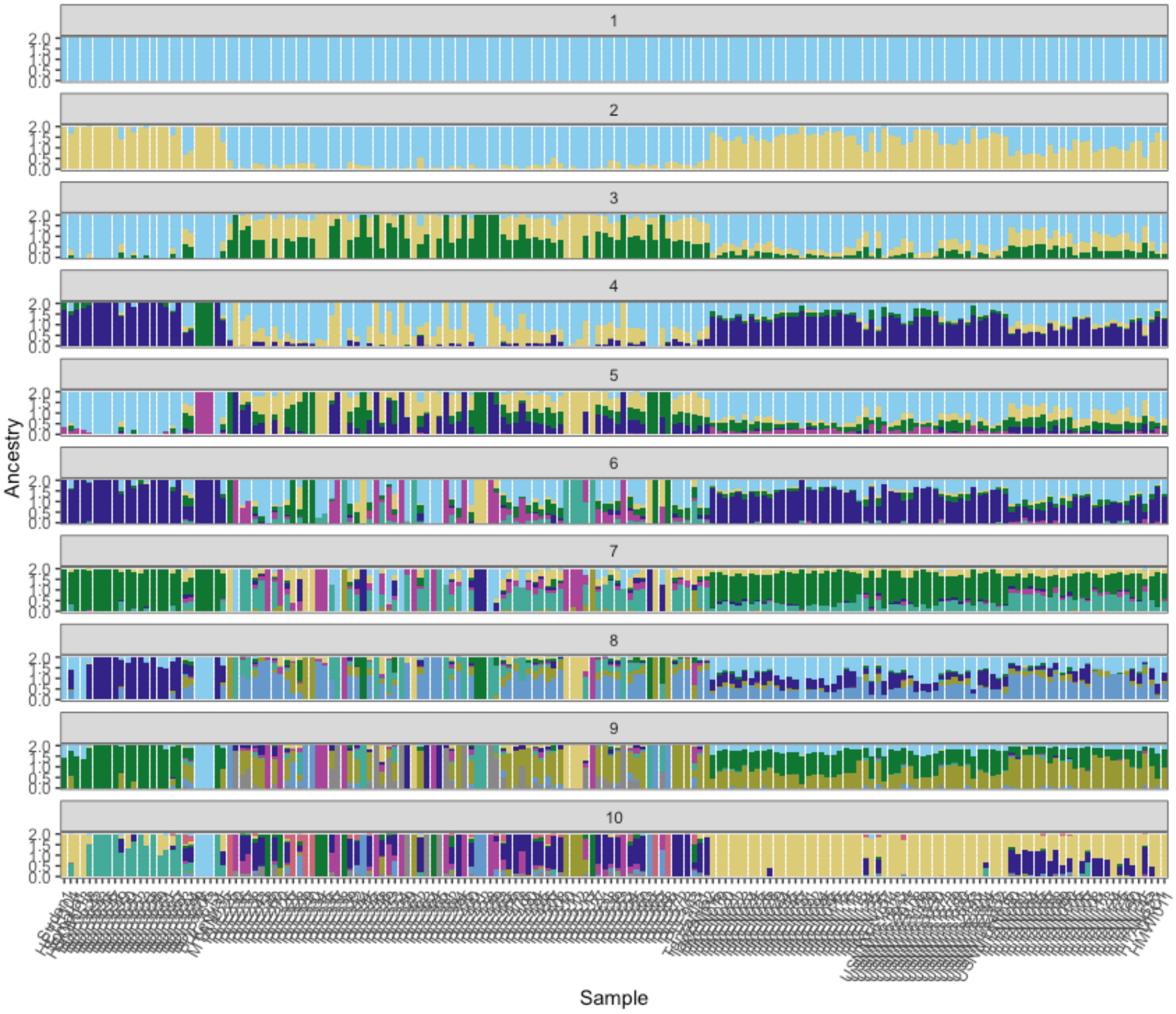


**Supplementary Figure S5**: NGSAdmix plots with K = 1-10, transitions removed, where samples had at least 1% of the genome covered.

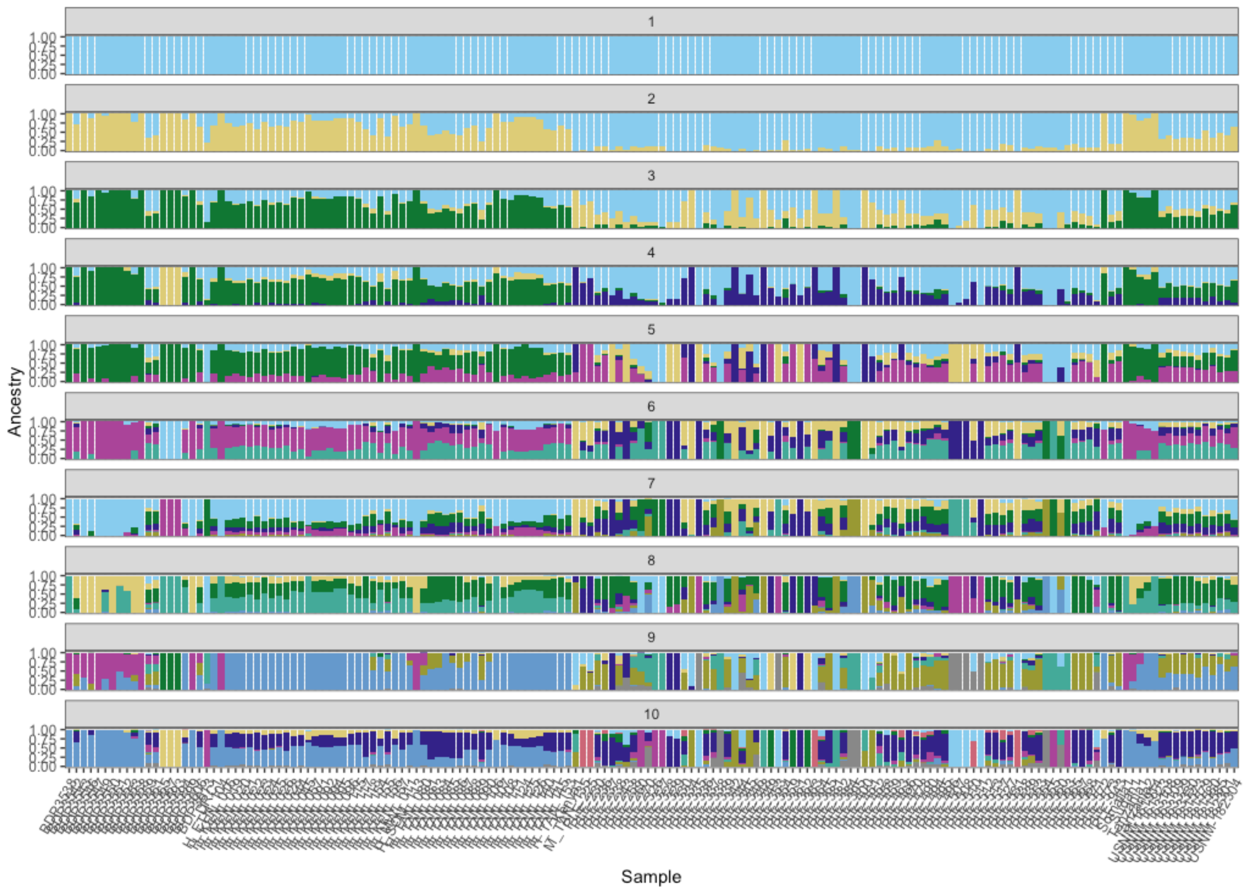


**Supplementary Figure S6**: NGSAdmix plots with K = 1-10, transitions removed, where samples had at least 5% of the genome covered.

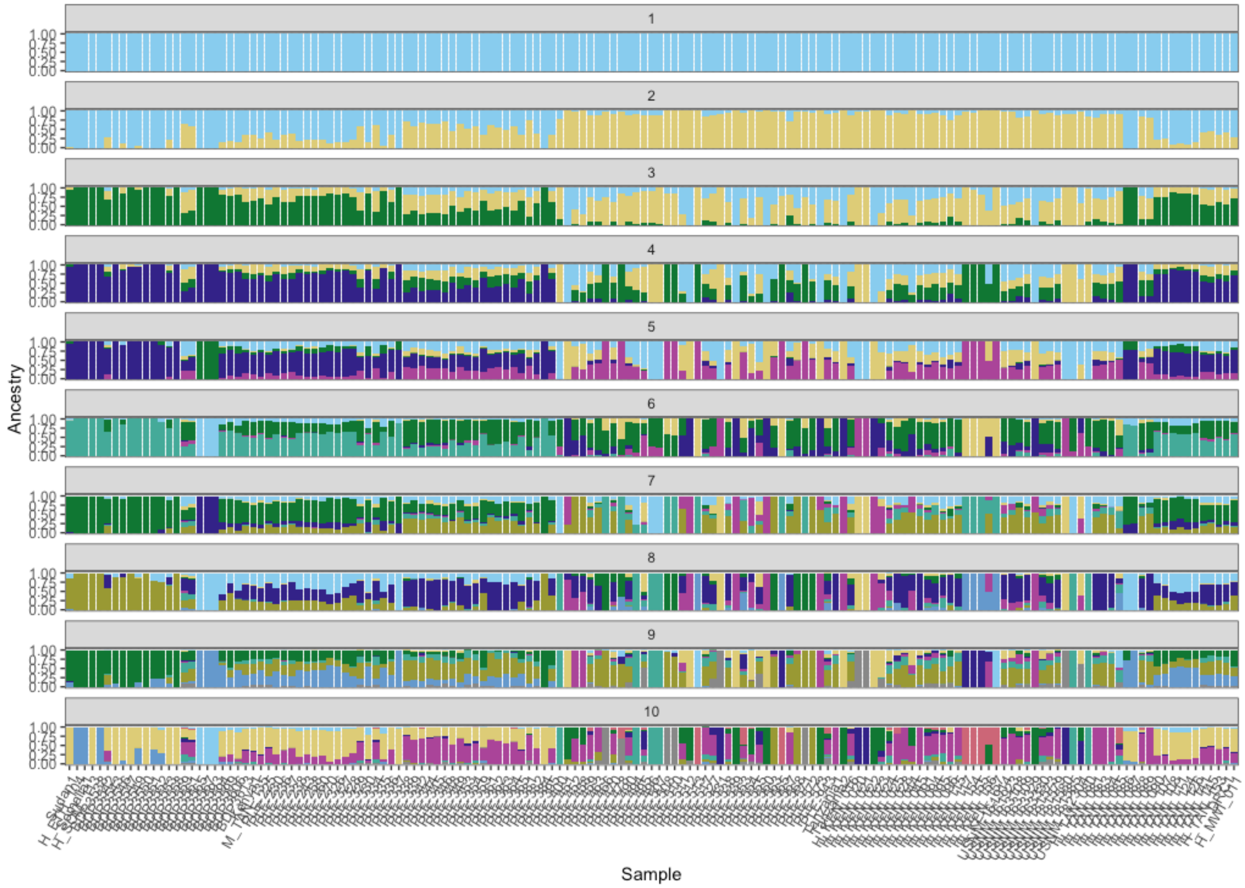


**Supplementary Figure S7**: NGSAdmix plots with K = 1-10, transitions removed, where samples had at least 10% of the genome covered.

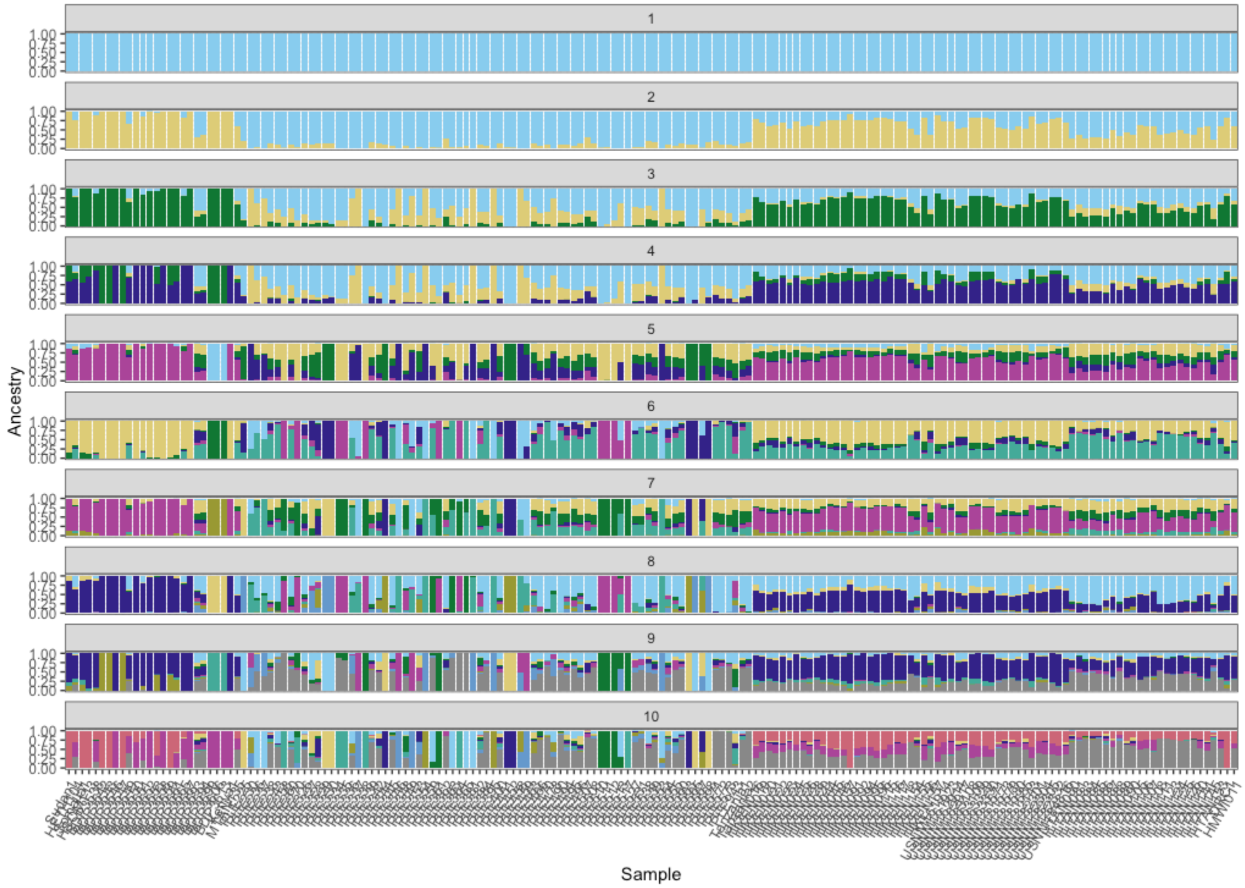


**Supplementary Figure S8**: NGSAdmix plots with K = 1-10, where samples had at least 1% of the genome covered.

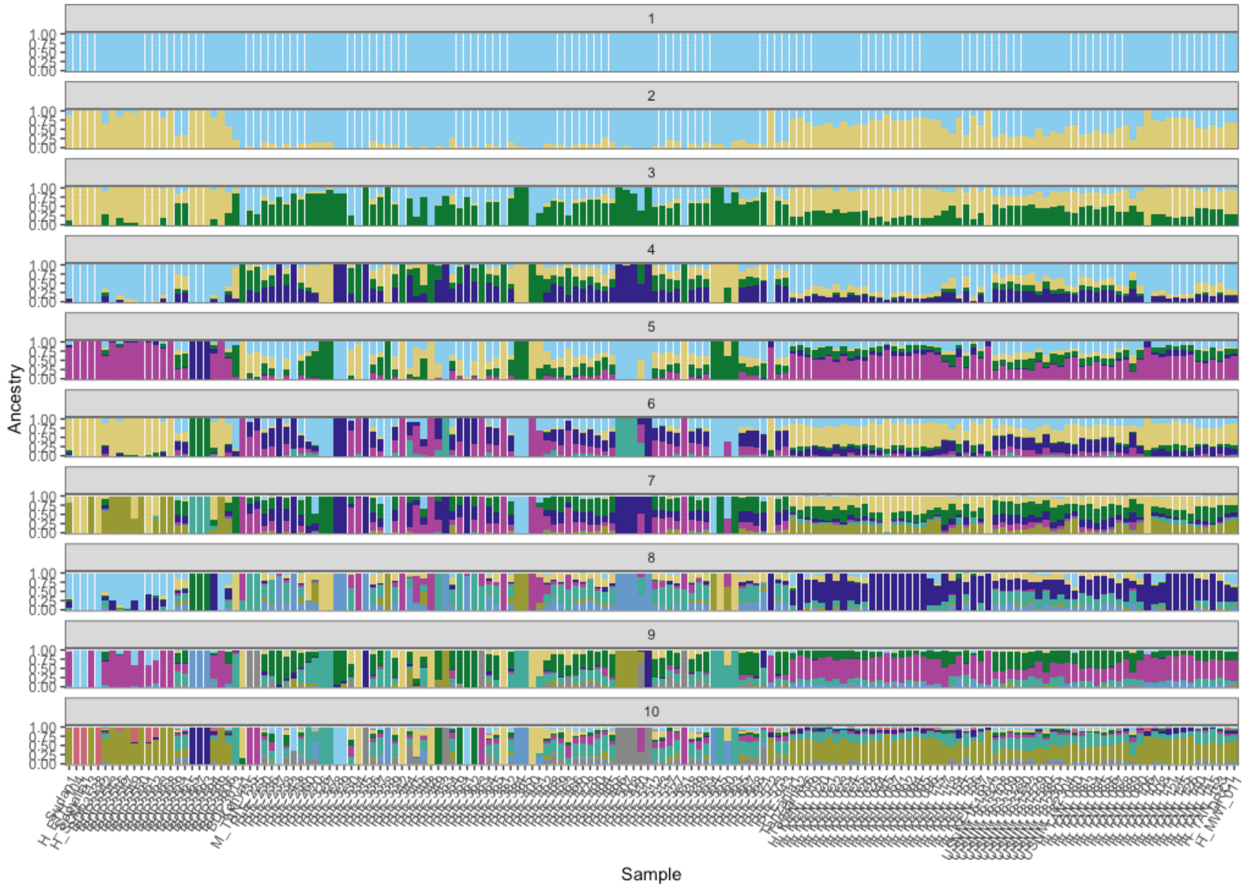


**Supplementary Figure S9**: NGSAdmix plots with K = 1-10, where samples had at least 5% of the genome covered.

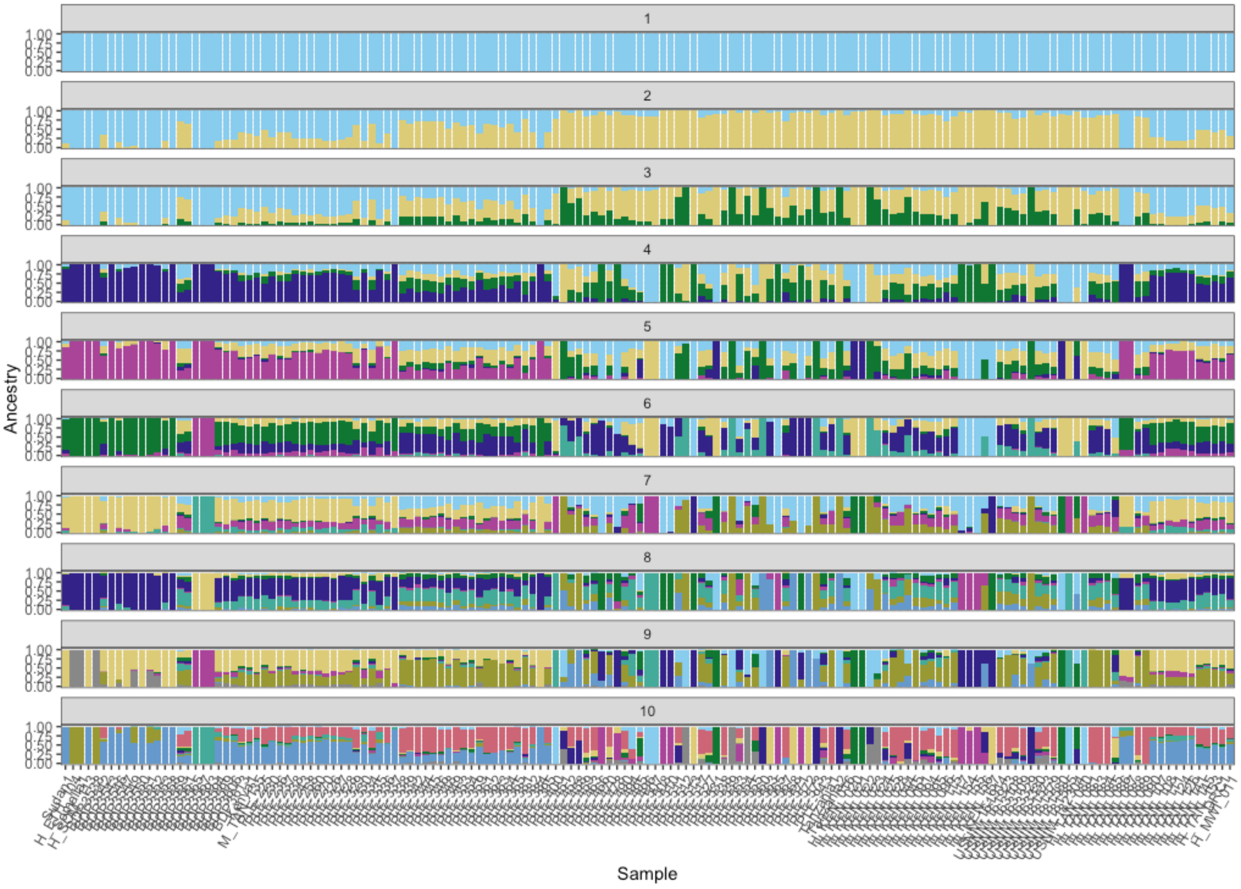


**Supplementary Figure S10**: NGSAdmix plots with K = 1-10, where samples had at least 10% of the genome covered.

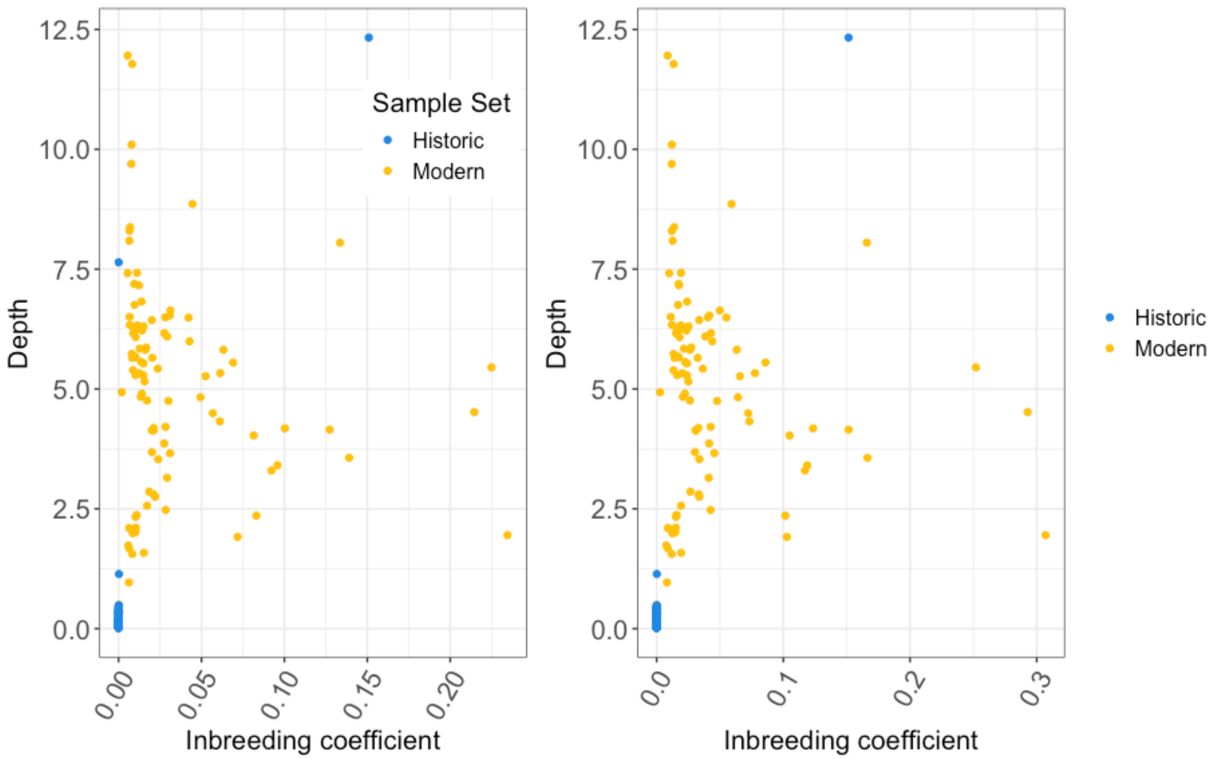


**Supplementary Figure S11**: Correlation of inbreeding and depth across samples (>1% coverage), split by estimates with no transition sites (left) and with transition sites included (right).

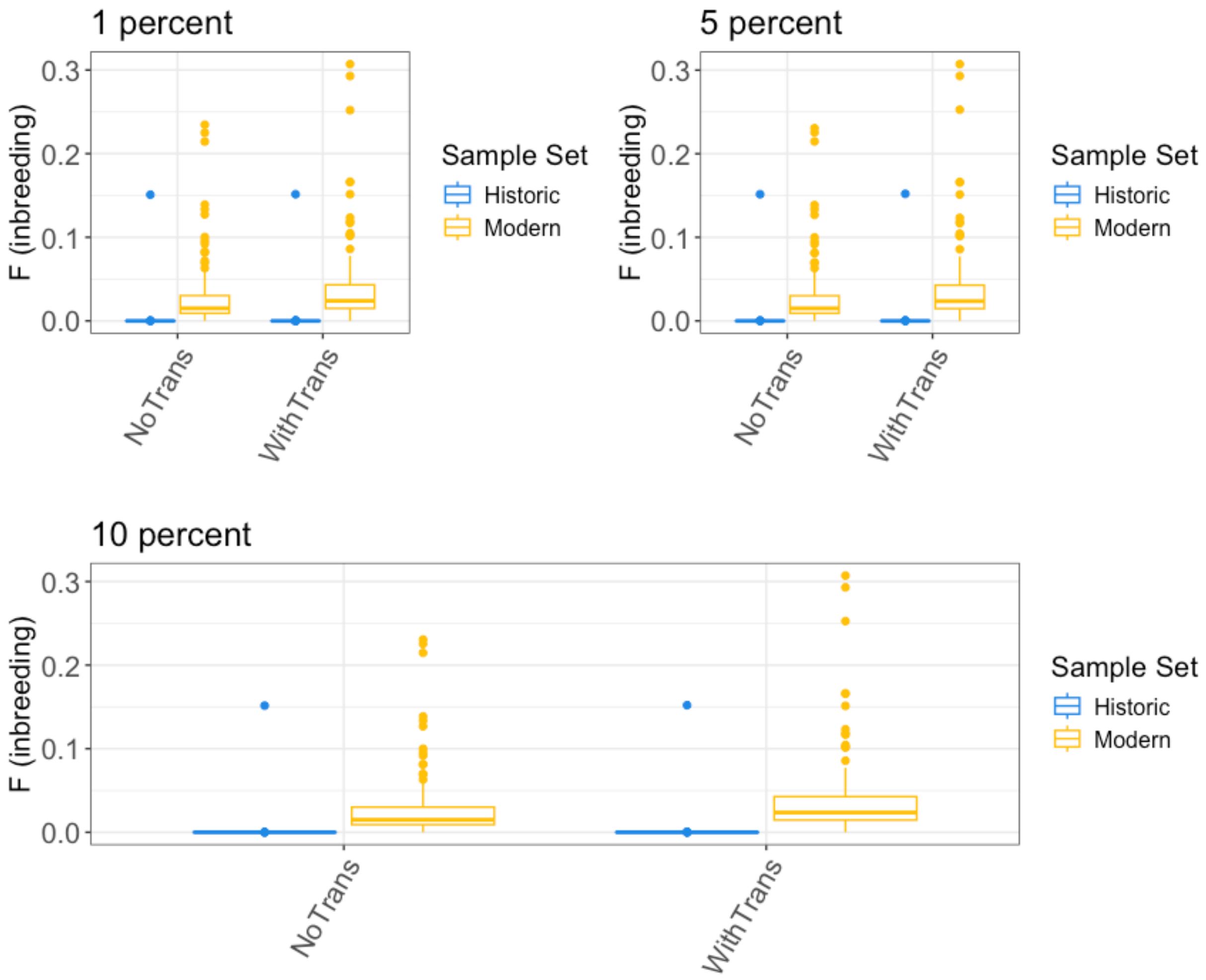


**Supplementary Figure S12**: Inbreeding estimates using ngsAdmix for recent and historic samples across various coverage minimums (1%, 5%, 10%), split by estimates with no transition sites included (left) and with transition sites included (right).

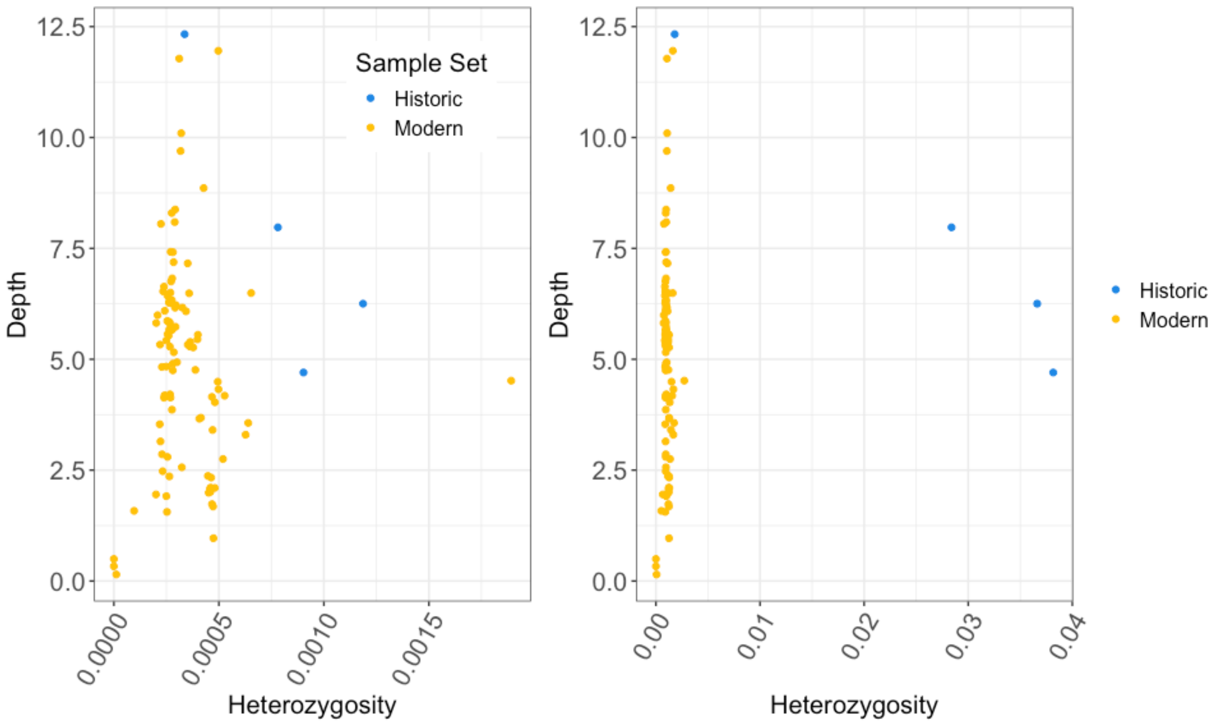


**Supplementary Figure S13**: Correlation of heterozygosity estimates with depth using ANGSD for recent (n = 100) and high coverage historic (n = 4) samples for transversions only (left panel) and all sites (right panel). Samples in the bottom left corner (FMNH 3559, FMNH 3550, and M_TAN_135) were removed from analyses due to low depth.

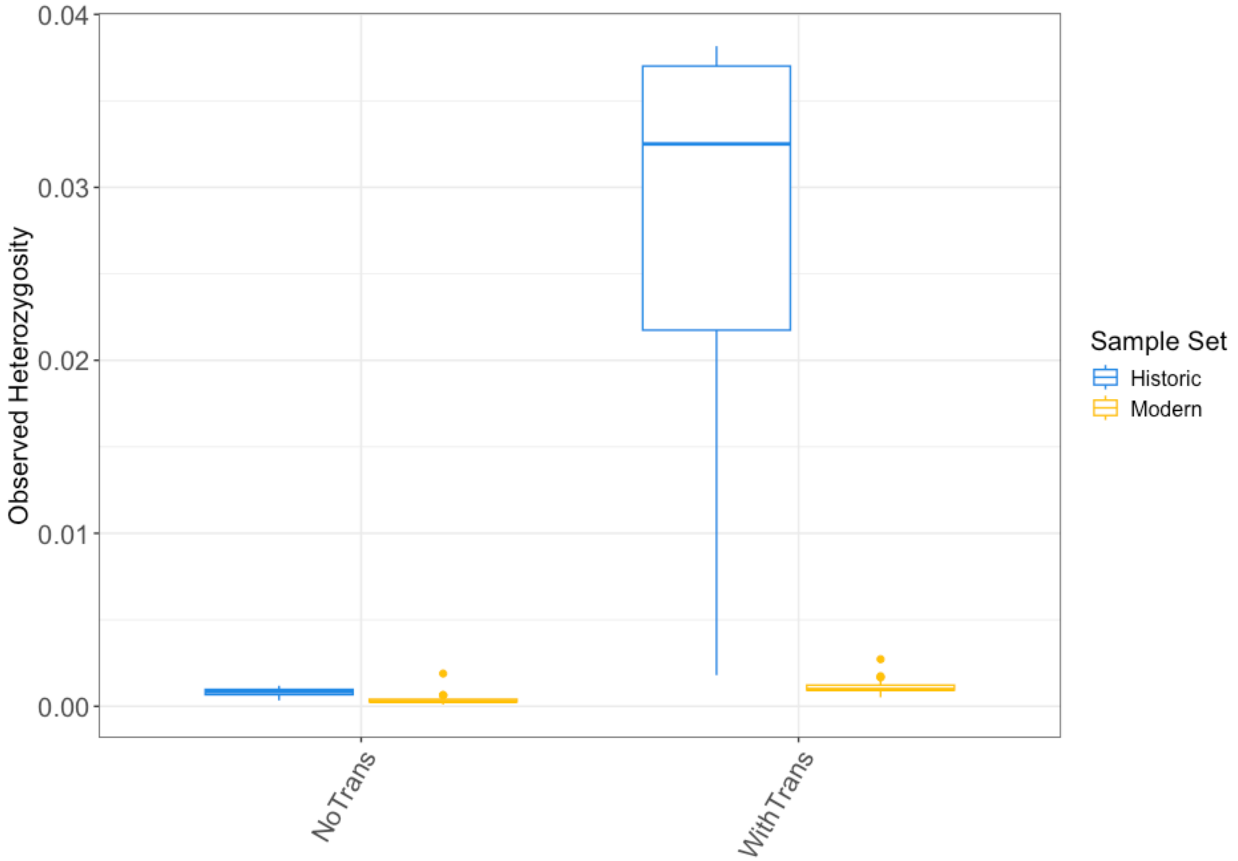


**Supplementary Figure S14**: Heterozygosity estimates using ANGSD for recent (n = 97) and high coverage historical (n = 4) samples, split by estimates with no transition sites included (left) and with transition sites included (right).

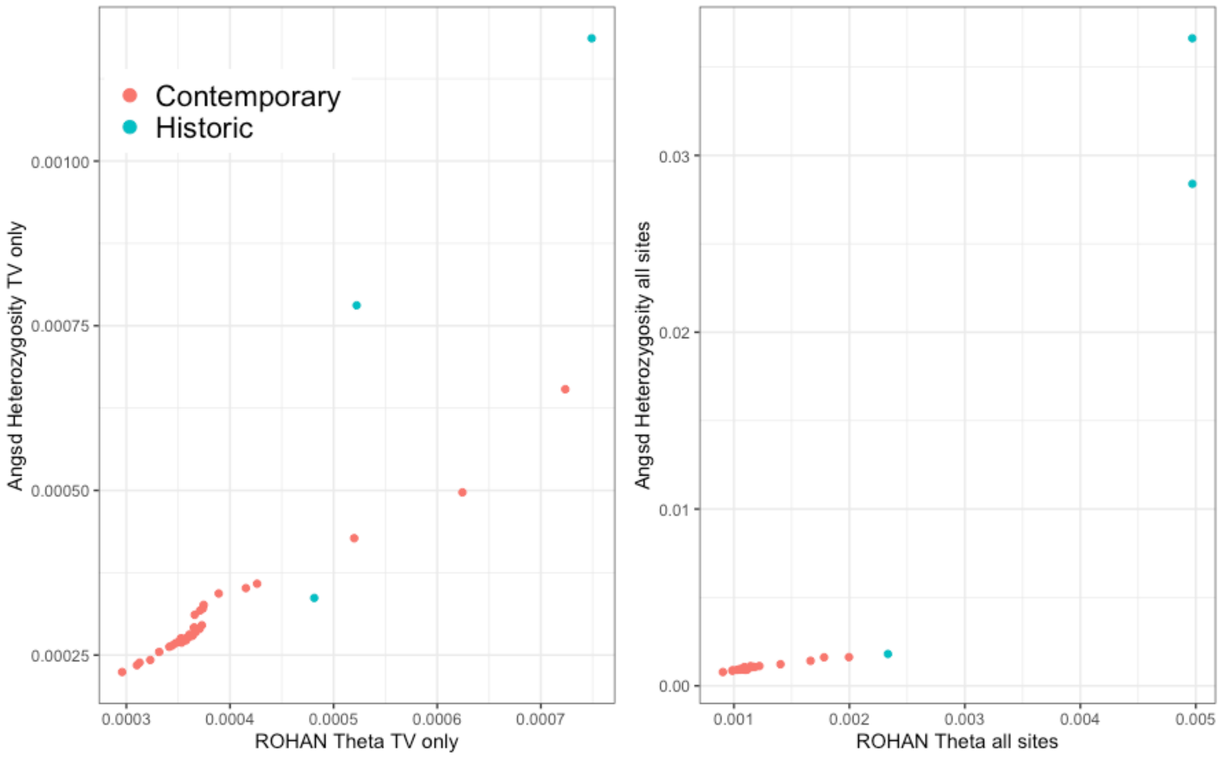


**Supplementary S15**: Comparison of theta estimates from ROHAN with observed heterozygosity estimates from ANGSD. Left panel shows only transversion sites, while the right panel shows all sites. Only individuals with an average depth of at least 6× included.

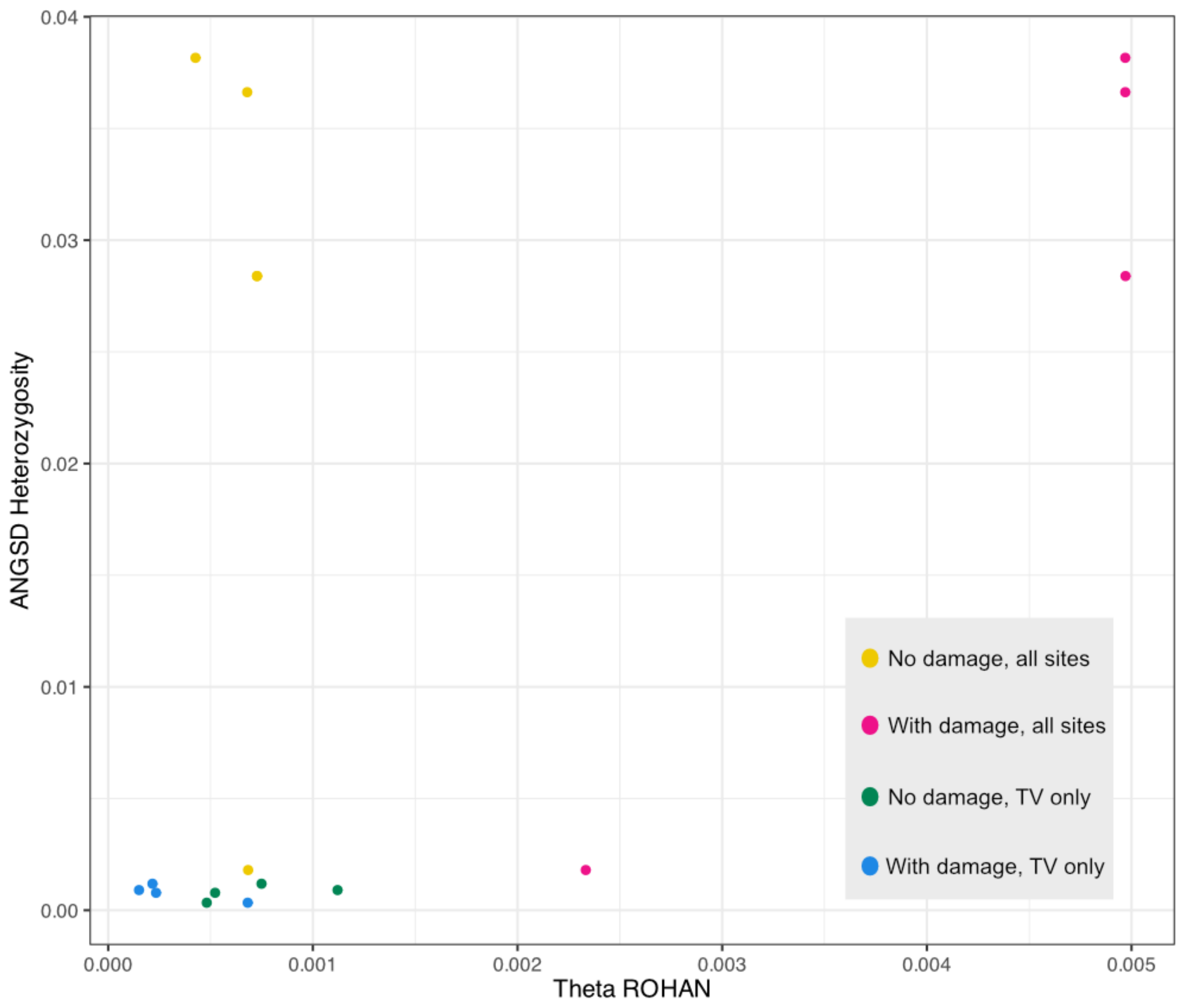


**Supplementary S16:** Comparison of various models of ROH calling using the historical data. Metric used for comparison is Theta (ROH inclusive) as output by ROHAN. TV only = transition sites removed.

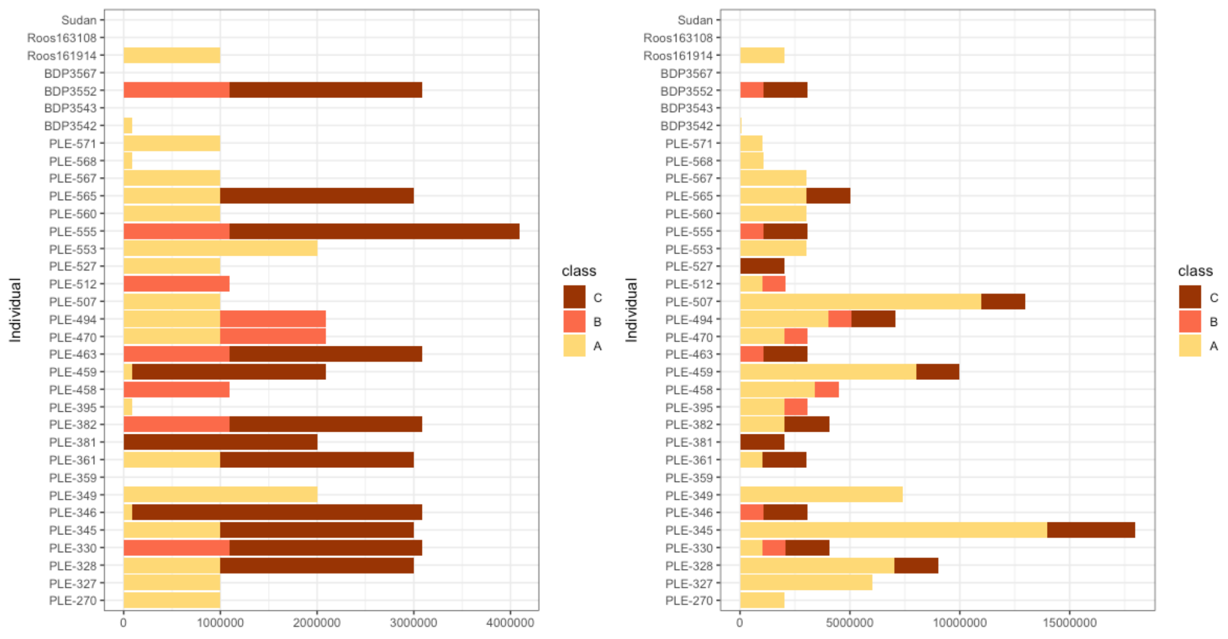


**Supplementary S17**: Estimates of ROH without (left) and with (left) transition sites removed across different size classes. We note the difference in the X axis. Historical samples are displayed at the top (Sudan, Roos163108, and 161914).
